# Pharmaco-Pheno-Multiomic Integration Reveals Biomarker Profiles and Therapeutic Response Prediction Models in Leukemia and Ovarian Cancer

**DOI:** 10.1101/2022.06.14.495846

**Authors:** Gilad Silberberg, Brandon Walling, Amy Wesa, Alessandra Audia, Ido Sloma, Yi Zeng, Guanghui Han, Jia Tang, Paige Pammer, A’ishah Bakayoko, Xuan Ren, Daniel Ciznadija, Bandana Vishwakarma, Yaron Mosesson, Marianna Zipeto, Michael Ritchie

## Abstract

Despite considerable progress made in improving therapeutic strategies, the overall survival for patients diagnosed with various cancer types remains low. Further, patients often cycle through multiple therapeutic options before finding an effective regimen for the specific malignancy being treated. A focus on building enhanced computational models, which prioritize therapeutic regimens based on a tumor’s complete molecular profile, will improve the patient experience and augment initial outcomes. In this study, we present an integrative analysis of multiple omic datasets coupled with phenotypic and therapeutic response profiles of Cytarabine from a cohort of primary AML tumors, and Olaparib from a cohort of Patient-Derived Xenograft (PDX) models of ovarian cancer. These analyses, termed <u>P</u>harmaco-<u>P</u>heno-<u>M</u>ulti<u>o</u>mic (PPMO) Integration, established novel complex biomarker profiles that were used to accurately predict prospective therapeutic response profiles in cohorts of newly profiled AML and ovarian tumors. Results from the computational analyses also provide new insights into disease etiology and the mechanisms of therapeutic resistance. Collectively, this study provides proof-of-concept in the use of PPMO to establish highly accurate predictive models of therapeutic response, and the power of leveraging this method to unveil cancer disease mechanisms.

## Introduction

In 2014, the Food and Drug Administration (FDA) issued regulatory guidance on the use of companion diagnostic (CDx) assays, which led to a rapid increase in the development of such assays. By the start of 2022, the total number of CDx assays approved by the FDA grew to 50 [1]. The majority of approved CDx assays are associated with therapeutics that treat hematologic and solid malignancies, reflecting a recent shift towards the development of targeted therapies in the treatment of cancer. Immunohistochemistry (IHC) and in situ hybridization (ISH) assays were the central technologies leveraged in a CDx assay until 2011, when the first polymerase chain reaction (PCR)-based CDx was approved. Since then, the use of PCR-based CDx assays has become a mainstay in the field, with 16 PCR-based assays currently approved [1]. More recently, next generation sequencing (NGS) has emerged as a common platform used in CDx assays under development. The first NGS assay to be approved by the FDA was the *FoundationFocus CDx*BRCA Assay in 2016 (Foundation Medicine). This assay detects deleterious alterations in BRCA1 and BRCA2 genes in patients with ovarian cancer and is used to assess whether a patient may be a candidate for treatment with Rucaparib (Rubraca). Since then, seven different NGS-based CDx assays have been approved by the FDA. Despite the increased adoption in the use of CDx platforms, all approved CDx assays are based on testing for the presence of one or two gene alterations as the basis for the prediction. Further, no CDx assay is available for non-targeted cytotoxic therapies, despite being dominant as front-line therapy options for most tumor indications [1]. Given the complex and dynamic interplay amongst dysfunctional biological pathways present in cancer cells, a more sophisticated approach to developing comprehensive CDx assays for all therapeutic strategies would increase prediction precision and broaden patient inclusion criteria.

Approximately 300,000 people are diagnosed with Acute Myeloid Leukemia (AML) each year. Although considerable progress has been made over the past 3 decades in developing successful treatments for AML, overall survival remains unacceptably low (< 30%) [2]. Several AML therapeutic treatment strategies exist, including an induction therapy regimen and a consolidation therapy regimen for those patients in remission. In patients under the age of 60, induction therapy often involves treatment with Cytarabine (ara-C) and an anthracycline drug such as Daunorubicin (Daunomycin) or Idarubicin. Despite promising responses achieved during induction therapy, only 40-50% of patients achieve complete response, with younger patients responding more favorably [3]. Currently, no approved CDx is available to help predict a patient’s response profile to Cytarabine. This is, in large part, because Cytarabine is a non-targeted cytotoxic agent, and no single gene aberration is linked to response profiles.

The overall survival of patients diagnosed with invasive epithelial ovarian cancer also remains low, with an average 5-year survival at less than 50% for all SEER stages combined [4]. The current initial treatment approach for patients with newly diagnosed advanced ovarian cancer includes the use of paclitaxel and carboplatin in combination with surgical cytoreduction. Remission is often achieved, but 80% of patients will have a recurrence within 3 years of remission [5]. Recent treatment advances include the use of poly(ADP)-ribose polymerase (PARP) inhibitors, such as Olaparib. The use of the *BRACAnalysis CDx* (Myriad Genetics) is employed to identify patients suitable for treatment with Olaparib. The basis for this CDx assay is the presence of deleterious alterations in the BRCA1 and BRCA2 genes as a criterion for the use of Olaparib. This effectively eliminates any patient with an unaltered BRCA1/BRCA2 gene status from the use of Olaparib, even though evidence exists to support the use of Olaparib in certain patients with wild-type (wt) BRCA status [6]. The development of a biomarker profile for PARP inhibitor response profiles, that broadens the molecular analysis, would expand inclusion criteria to a wider audience of patients that could benefit from treatment.

In this study, we provide proof-of-concept for an integrative approach to building complex computational models that incorporate multiple omic datasets to reconstruct tumor cell biology and predict response/resistance profiles more broadly. We performed this analysis, termed PPMO, for non-targeted (Cytarabine) and targeted (Olaparib) therapeutics, and in extremely heterogenous hematologic (AML) and solid (Ovarian) malignancies. The PPMO-based prediction models were able to accurately predict prospective therapeutic response profiles for subsequently investigated tumors. Further, the PPMO models provided supportive evidence of previously identified biomarkers and therapeutic targets and highlight several potentially tractable and previously unreported therapeutic targets.

## Results

### AML Patient Characteristics

Primary samples (derived from a leukapheresis procedure) from 23 patients diagnosed with AML were collected. The patients presented with a wide distribution of AML model classes, as determined by the French-American-British (FAB) classification, with forty three percent of patients being treatment naïve (Fig. 1a-b, Supplementary Table 1). A wide distribution of age and disease status was also observed amongst the cohort of patients (Fig. 1c-d, Supplementary Table 1). Pathogenic mutations in *IDH1/2, FLT3, NPM1, CUX1, RUNX1, KIT, DNMT3A, STAG2, and TP53* were represented across the patient landscape, highlighting molecular diversity of this cohort (Supplementary Table 1). A similarity network fusion (SNF) clustering analysis of RNA and protein expression revealed 4 clusters within the patient cohort, with clusters 1 & 2 exhibiting largely abnormal cytogenetics. No other major discriminating factors were easily observed between the clusters generated from this SNF analysis, demonstrating molecular heterogeneity and diversity amongst all tumors within this cohort (Fig. 1e). Further, the superficial SNF did not establish a cluster association with cytarabine response profiles. Collectively, these results demonstrate a diversity in this cohort of tumor samples and highlights the need for complex molecular and phenotypic analysis to reveal the underpinnings of cytarabine resistance and build a response biomarker profile.

**Fig.1:**
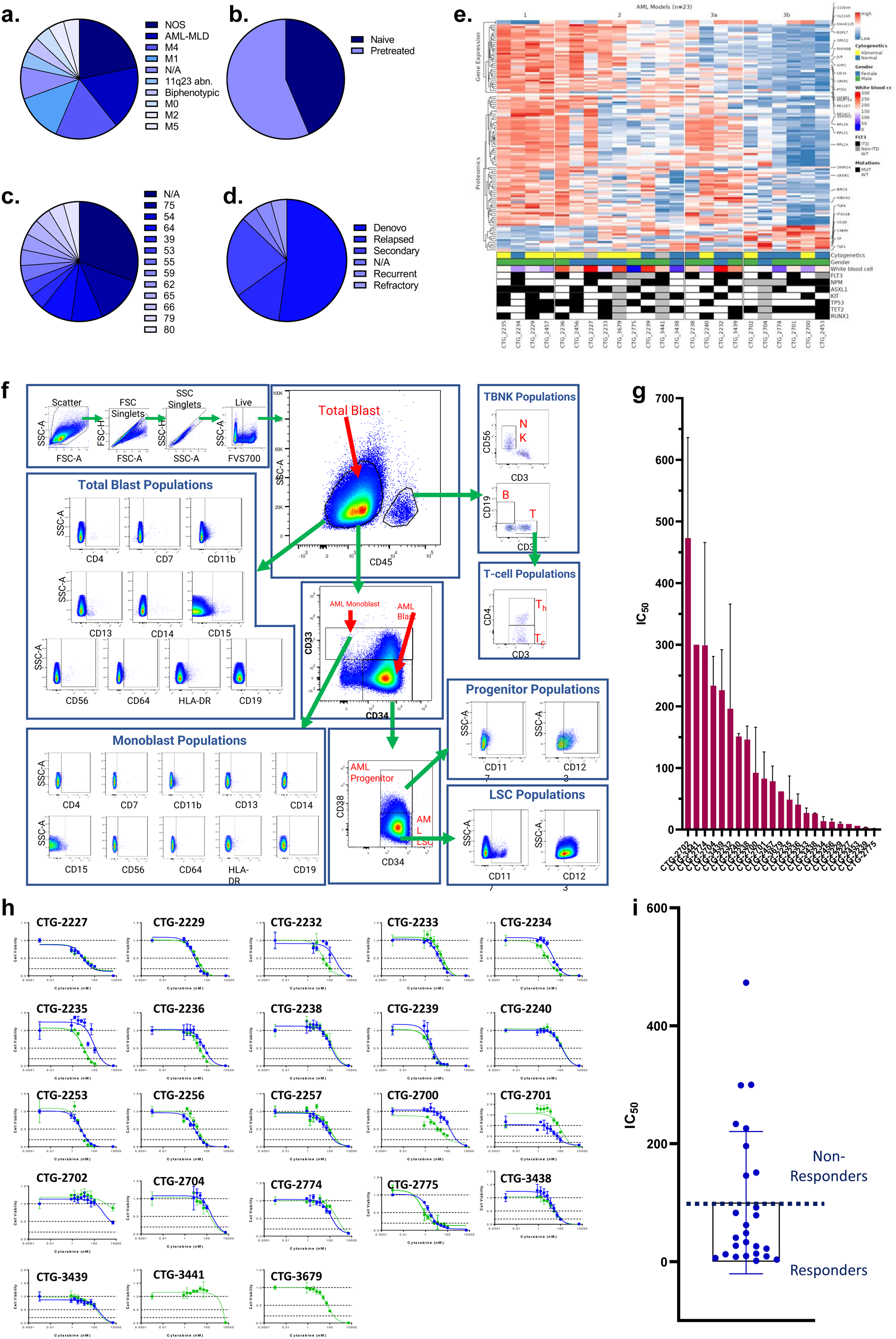
Clinical characteristic of the patients and primary AML samples are denoted, including **(a)** the distribution of FAB classes across all samples, **(b)** the distribution of treatment naïve and pretreated tumors, **(c)** the age distribution of all patients the samples were collected from, and **(d)** the disease status of the primary samples. (**e)** Similarity network fusion (SNF) analysis of combined RNA (upper heatmap panel) and protein expression (lower heatmap panel) data obtained from AML samples **(f)** representative example of the flow cytometry gating strategy for *PhenoSeek*. **(g)** *ex vivo-based* IC_50_ values for cytarabine treatment were acquired in primary AML samples and plotted as a mean with SEM for each sample. **(h)** The acquisition of Cytarabine-mediated IC_50_ curves across all patients’ samples are shown. For instances where two different IC50 values were acquired, different tests are shown in green or blue. **(i)** The mean IC_50_ value of Cytarabine for each sample is plotted and models are binarized as responders or non-responders based on an IC_50_ cutoff of 100 nM.

### AML Phenotyping

Given the complex heterogeneity of cell populations observed amongst different AML tumors, the AML samples were first subjected to extensive cell diversity phenotyping via flow cytometry. This included a comprehensive AML-specific 19-channel flow cytometry panel performed under Good Clinical Laboratory Practice (GCLP) guidelines (called *PhenoSeek*) (Supplementary Table 2 and 3). The gating strategy for this flow panel enables the identification of 35 different cellular subsets (Fig. 1f). A diversity of cell populations across all patients are observed, and a representative diversity of a select number of cell populations is presented in Supplementary Fig. 1a-c.

The primary AML samples were also subjected to WES, RNAseq, quantitative(Q)-proteomics for protein expression, and Phosphorylation(P)-proteomics to establish the phosphorylation state of each protein. Kinase activity within the tumors was established using a <u>s</u>ingle <u>s</u>ample <u>G</u>ene <u>S</u>et <u>E</u>nrichment <u>A</u>nalysis (ssGSEA). Importantly, we find only a 17.3% correlation between RNA expression and protein expression amongst these tumor samples (Supplementary Fig.2a). Further, we find a very low correlation between protein expression and kinase activity, reinforcing the different post-translational processes that regulate protein activity (Supplementary Fig. 2b). Collectively, these results highlight the advantage to multiomic integration analysis when using omics to reconstruct a tumor’s molecular makeup, and the shortcomings of relying on only RNA expression or protein expression to understand protein or pathway activity.

These primary samples were also subjected to a primary *ex vivo* cytotoxicity assay to establish sensitivity and resistance status to cytarabine. The *ex vivo* protocol subjects the patient samples to short term culture (<10 days), to ensure lack of genetic drift, and clonal selection within the samples. This type of primary explant culture is necessary due to the propensity for AML samples to rapidly differentiate and clonally select[7-9]. The assay demonstrated strong reproducibility across the models tested and provided a suitable range of sensitivity to cytarabine treatment ((Fig. 1g-h). Cytarabine responses measured in the patient samples were binarized into ‘responder’ (R) and ‘non-responder’ (NR) categories using an IC_50_ cutoff of 100 nM (Fig.1i).

### Integrated Phenotype & Omics Analysis of Cytarabine Response in Primary AML

To study the molecular mechanism underlying cytarabine response and identify candidate biomarkers, *Data Integration Analysis for Biomarker discovery using Latent Components* (DIABLO)[10], based on partial least squares discriminant analysis (PLS-DA), was applied on all omics blocks (Supplementary Fig.3a-b). We first performed tuning steps to assess the required number of latent variables, and the optimal number of features to be selected from each block in each latent variable (see Materials and Methods). Using these parameters, 2 latent variables were calculated for each block. We observed that, as expected, latent variables of different blocks tended to co-cluster by components (Fig. 2a). Importantly, Component 1 (Comp 1-Cytarabine) was strongly correlated with cytarabine sensitivity, (r = 0.519 kinase block, 0.702 mutation block, 0.822 protein block, and 0.78 RNA block (Supplementary Fig. 3c)). Hierarchical clustering of all Comp 1-Cytarabine features differentiates response groups clearly (Fig.2b).

**Fig.2:**
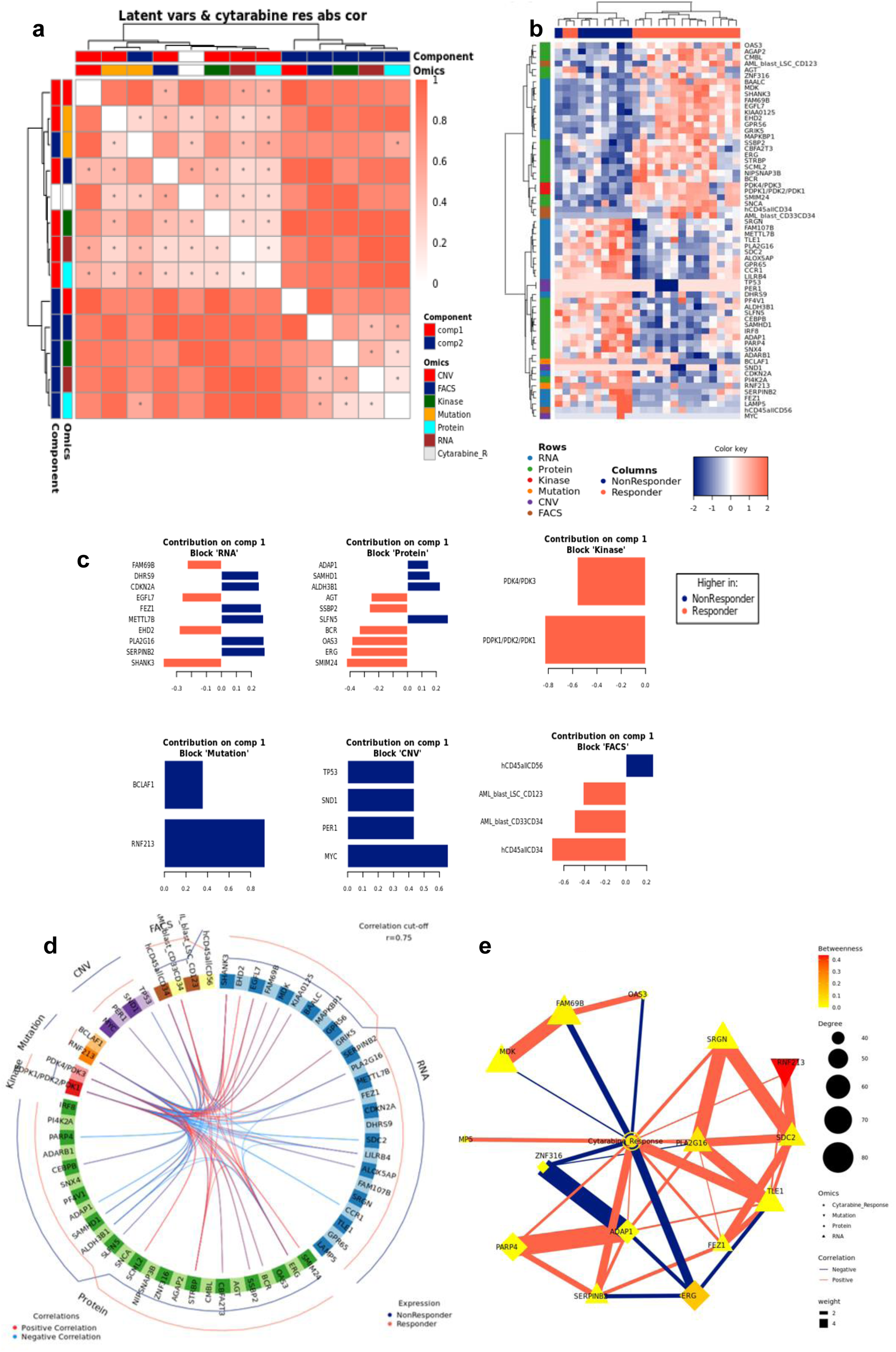
(a) Pairwise correlations between latent variables (components) calculated for the different omics blocks, and cytarabine response (shown in white in the row/column legends). Dots indicate significant correlations (FDR < 0.1, see fig. S7C). The first component of all blocks is strongly correlated with cytarabine response. (b) Clustering of omic blocks contained within Comp 1-Cytarabine. (c) Features with the highest contribution to component 1 of each omics block. (d) Top correlations (r >= 0.8 / r <= -0.8) between features contributing to component 1, between and within omics blocks (inside the circle). Lines outside the circle indicate the features expression level for the responders and non-responder’s groups, where distance from the circle corresponds to expression level. (e) A Context Likelihood or Relatedness network (CLR) calculated over multi-omics features from Comp 1-Cytarabine, along with cytarabine response IC50. Shown here is a 1-neighborhood subgraph around cytarabine response, filtered for strongest (edge weight > 2) and significant (correlation FDR < 0.05) relationships. Edge thickness indicates interaction strength (weight).

The loadings of each feature on Comp 1-Cytarabine also indicate the strength of their relationships with cytarabine response. Features with the highest absolute loadings in their blocks include AML blasts (CD34+) with higher levels in R, CD56+ cells with higher levels in NR, CNVs in MYC (more frequent in NR), mutant RNF213 (more frequent in NR), Pyruvate Dehydrogenase Kinase (PDK)-family kinase activity (higher in R), SAM domain and HD domain-containing protein 1 (SAMHD1) protein (higher in NR), Serpin Family B Member 2 (SERPINB2) RNA (higher in NR), Methyltransferase-like 7B (METTL7B) RNA (higher in NR), and Small Integral Membrane Protein 24 (SIMM24) Protein (higher in NR), among others (Fig. 2c). In most omics blocks, the first 2 variates were able to discriminate well between the response classes, where the proteomics block variate 1 excelled (Supplementary Fig. 3a). A Functional enrichment analysis calculated for proteins and transcripts ranked by their absolute Comp1-cytarabine loadings revealed enrichment in fatty acid metabolism, among others (Supplementary Fig. 3d).

The strongest correlations between Comp 1-cytarabine features in R-associated tumors revealed positive correlations between CD34+ cell enrichment and several features, including higher kinase activity of PDK family of proteins, ETS-Related Gene (ERG) protein expression, and Epithelial growth factor-like 7 (EGFL7) RNA expression (Fig. 2d). In NR-associated tumors, strong negative correlations between CD34+ cell populations and SAMHD1 protein expression, Aldehyde dehydrogenase 3B1 (ALDH3B1) protein expression, CCAAT enhancer binding protein gamma (CEBPB) protein expression, and leukocyte immunoglobulin-like receptor subfamily B4 (LILRB4) RNA expression was observed (Fig. 2d). Strong positive correlations between mutations in RNF213 and poly(ADP-ribose) polymerase 4 (PARP4) protein expression and LILRB4 RNA expression was also observed in tumors with NR cytarabine profiles (Fig. 2d). Context Likelihood or Relatedness (CLR) was used to reconstruct the regulatory network over multi-omics features from Comp 1-Cyarabine, along with cytarabine response (IC_50_), based on mutual information (Fig. 2e). This network represents the pan-omics molecular signature underlying/determining cytarabine response predisposition, as well as the interplay among them. The most strongly linked direct neighbors of the cytarabine response profile include known oncogenes and prognostic markers of AML, including Midkine (MDK) RNA [11], ERG protein [12], SERPINB2 RNA [13], and Serglycan (SRGN) RNA [14], among others (Fig. 2e). The 2-neighborhood subgraph including a more extensive features network (Fig. S4) features several additional known oncogenes and prognostic markers for AML, including METTL7B RNA [15], LILRB4 RNA [16], SAMHD1 protein [17, 18] and EGFR7 [19].

### Ovarian Patient Characteristics and Olaparib Profiling

Primary ovarian samples (derived from biopsy or surgical resection) from 24 patients diagnosed with ovarian cancer were collected and used to establish Patient-Derived Xenograft (PDX) models (Supplementary table 5). Most samples were established from metastatic lesions (Fig. 3a) and from a wide distribution of patient ages (Fig.3b). 16 of the patients were pretreated with platinum-based therapy regimens (Fig.3c) and most were of an advanced stage (Fig. 3d). The PDX models were also subjected to WES, RNAseq, Q-proteomics and P-proteomics. Kinase activity within the tumors was established using ssGSEA. In contrast to the AML cohort, we find a higher correlation (86%) between RNA expression and protein expression amongst these tumor samples (Supplementary Fig.5a). However, we find a very low correlation between protein expression and kinase activity (Supplementary Fig.5b)

**Fig.3:**
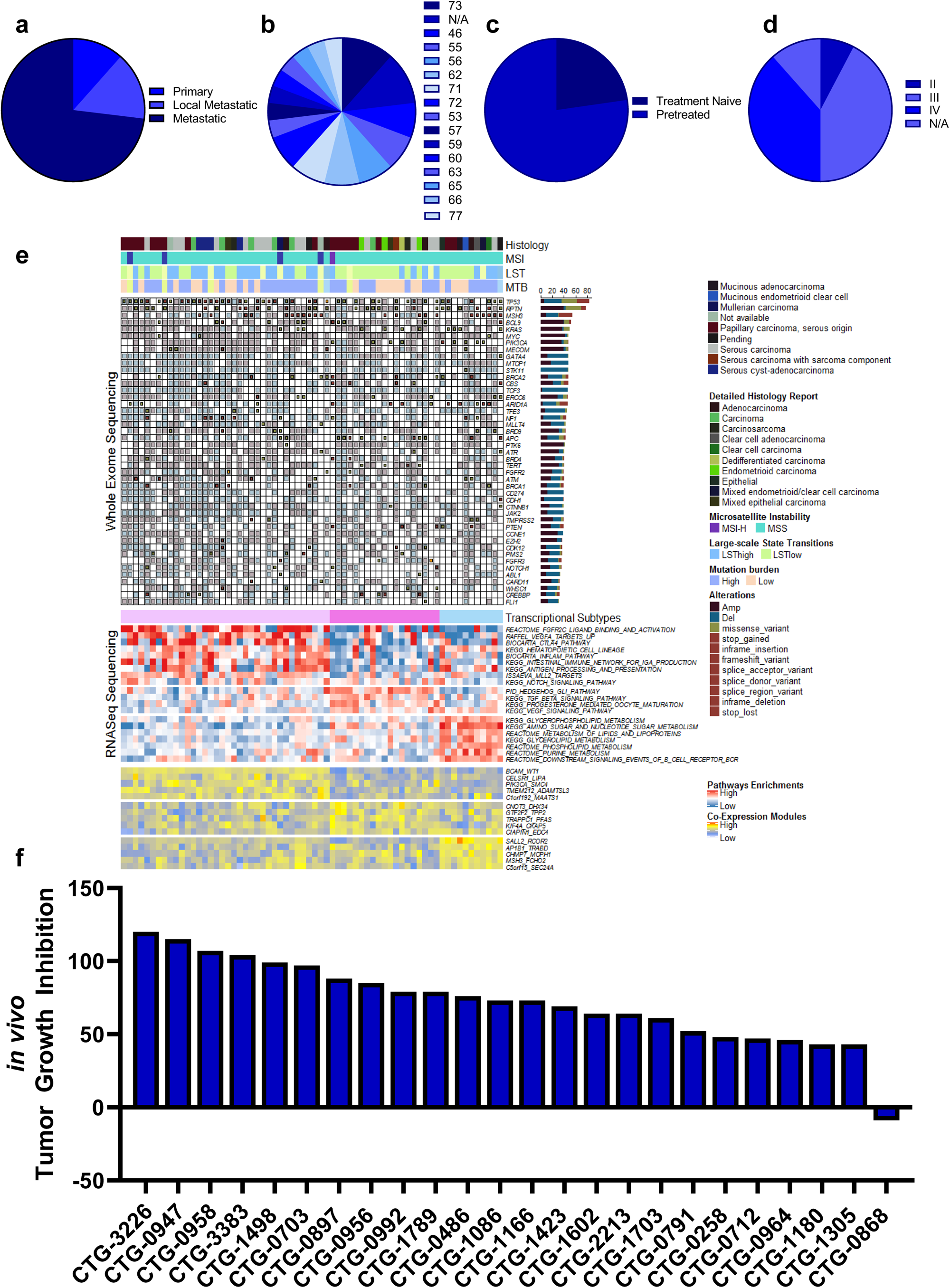
Clinical characteristic of the patients and ovarian PDX models are denoted, including (a) primary or metastatic status, (b) the age distribution of all patients the samples were collected from, (c) the distribution of treatment naïve and pretreated tumors, and (d) the disease stage of the primary tumors. (e) Similarity network fusion (SNF) analysis of combined RNA expression (upper heatmap panel) and gene set enrichment (lower heatmap panel) data obtained from the PDX models, with additional information on mutation status shown at the top. (f) in vivo-based TGI values for Olaparib treatment were acquired in ovarian PDX models and plotted as a waterfall plot.

A similarity network fusion (SNF) clustering analysis of RNA and protein expression revealed 3 major transcriptional subtypes within this ovarian cancer cohort (Fig. 3e). BRCA1 and BRCA2 mutational status was represented broadly across all three transcriptional subtypes, with no other phenotypic characteristic associating strongly with any of the clusters (Fig.3e). These PDX models were also subjected to a subcutaneous *in vivo* efficacy to establish sensitivity and resistance status to Olaparib. Tumor growth inhibition (TGI) measurements related to Olaparib treatment fell within a range of response, providing an effective dataset for a PPMO integration (Fig.3f).

### Integrated Omics Analysis of Olaparib Response in Ovarian PDX

To study the molecular mechanism underlying Olaparib response and identify candidate biomarkers, PPMO analysis was again employed. Since we obtained a range of *in vivo* responses to Olaparib without a clear cutoff between response extremes, we integrated Olaparib pharmacology as a regression analysis, rather than a binarized response set. This also enabled us to test an alternative approach to PPMO integration, whereby we were attempting to predict specific TGI values.

We first performed tuning steps to assess the required number of latent variables, and the optimal number of features to be selected from each block in each latent variable (see Materials and Methods). Using these parameters, 5 latent variables were calculated for each block. We observed latent variables of different blocks co-clustering by components in this analysis (Fig. 4a). Importantly, Component 1 in this analysis (Comp 1-Olaparib) was strongly correlated with Olaparib sensitivity. Hierarchical clustering of all Comp 1-Olaparib features differentiates response groups clearly (Fig.4b). The loadings of each feature on Comp 1-Olaparib also indicate the strength of their relationships with Olaparib response. Features with the highest absolute loadings in their blocks include kinase activity of MOS Proto-Oncogene, Serine/Threonine Kinase (MOS), Citron Rho-Interacting Serine/Threonine Kinase (CIT) protein expression, Endothelial Cell Adhesion Molecule (ESAM) RNA expression, mutations in KIAA1549 and CNVs in Cadherin 11 (CDH11) associating more with tumors on the non-responding side of the regression analysis (Fig.4c). Features with the highest absolute loadings associating more with tumors on the responding side of the regression analysis include kinase activity of Serine/Threonine-Protein Kinase BRSK2 (BRSK2), kinase activity of Protein Kinase AMP-Activated Catalytic Subunit Alpha 2 (PRKAA2), protein expression of Asparaginase And Isoaspartyl Peptidase 1 (ASRGL1), protein expression of Leucine Rich Repeats And Immunoglobulin Like Domains 1 (LR1G1), RNA expression of G Protein-Coupled Receptor 157 (GPR157), and CNV in Beta-2-Microglobulin (B2M), among others. (Fig.4c). A Functional enrichment analysis calculated for proteins and transcripts ranked by their absolute Comp1-Olaparib loadings revealed enrichment in Serine/Threonine kinase activity and Cadherin-related networks in tumors associating with poor response to Olaparib in the regression analysis (Supplementary Fig. 6).

**Fig.4:**
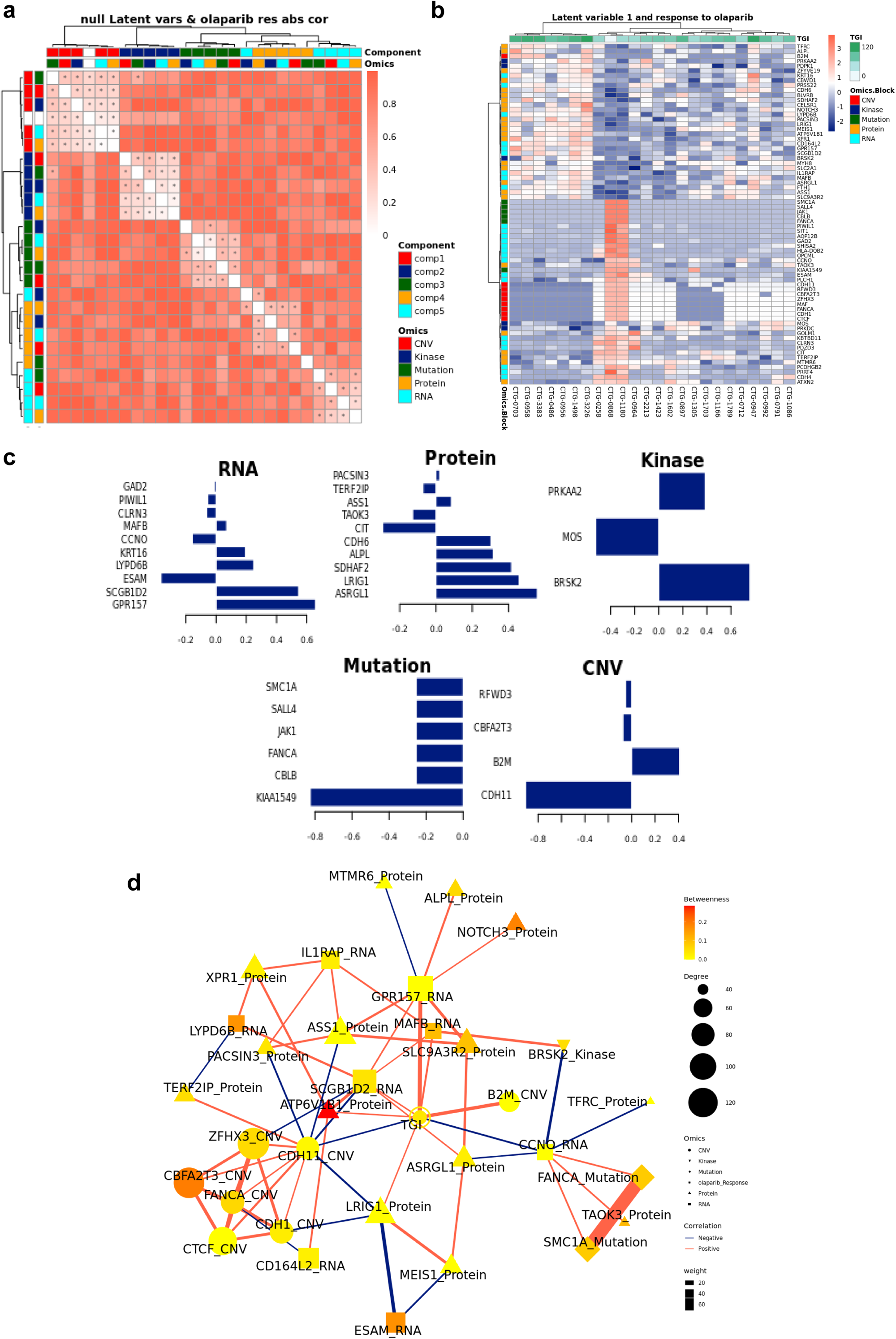
(a) Pairwise correlations between latent variables (components) calculated for the different omics blocks, and olaparib response (shown in white in the row/column legends). Dots indicate significant correlations (FDR < 0.1, see fig. S7C). The first component of all blocks is strongly correlated with olaparib response. (b) Clustering of omic blocks contained within Comp 1-Olaparib. (c) Features with the highest contribution to Comp 1-Olaparib of each omics block. (d) A Context Likelihood or Relatedness network (CLR) calculated over multi-omics features from component 1, along with olaparib response TGI Shown here is a 2-neighborhood subgraph around Olaparib response, filtered for strongest (edge weight > 2) and significant (correlation FDR < 0.05) relationships. Edge thickness indicates interaction strength (weight).

Context Likelihood or Relatedness (CLR) was used to reconstruct the regulatory network over multi-omics features from Comp 1-Olaparib, along with Olaparib response (TGI), based on mutual information (Fig. 4d). The most strongly linked direct neighbors of the Olaparib response profile include known oncogenes and prognostic markers of ovarian cancer, including Xenotropic And Polytropic Retrovirus Receptor 1 (XPR1) protein expression [20], Notch Receptor 3 (NOTCH3) protein expression [21], MAF BZIP Transcription Factor B (MAFB) RNA expression [22], Argininosuccinate synthetase 1 (ASS1) protein expression [23], and MEIS1 protein expression [24], among others. (Fig. 2e).

### Prospective Prediction of Cytarabine and Olaparib Responses

Though there were numerous biomarkers highlighted in our Cytarabine and Olaparib PPMO-based models, no one of them is strong enough to serve as an independent biomarker predicting therapeutic sensitivity. We therefore sought to test the trained PPMO models as a basis for prospective therapeutic sensitivity prediction. The resulting Comp 1-Cytarabine-based model was tested on 7 newly profiled AML samples, which had molecular and response data, but lacked FACS data. Despite the lack of the FACS block, the computational model was able to correctly classify the therapeutic response profile of 6 samples (Fig. 5a-b). Importantly, removing any omic block within the PPMO model led to a significant reduction in the predictive power, reinforcing the notion that single gene or single omic-block biomarkers have limited potential. Notably, the mis-classified sample was on the border between an R and NR threshold (IC_50_ of 100 nm), indicating that there may be a ‘gray zone’ within the model whereby tumors will be difficult to predict a given response profile.

**Figure 5:**
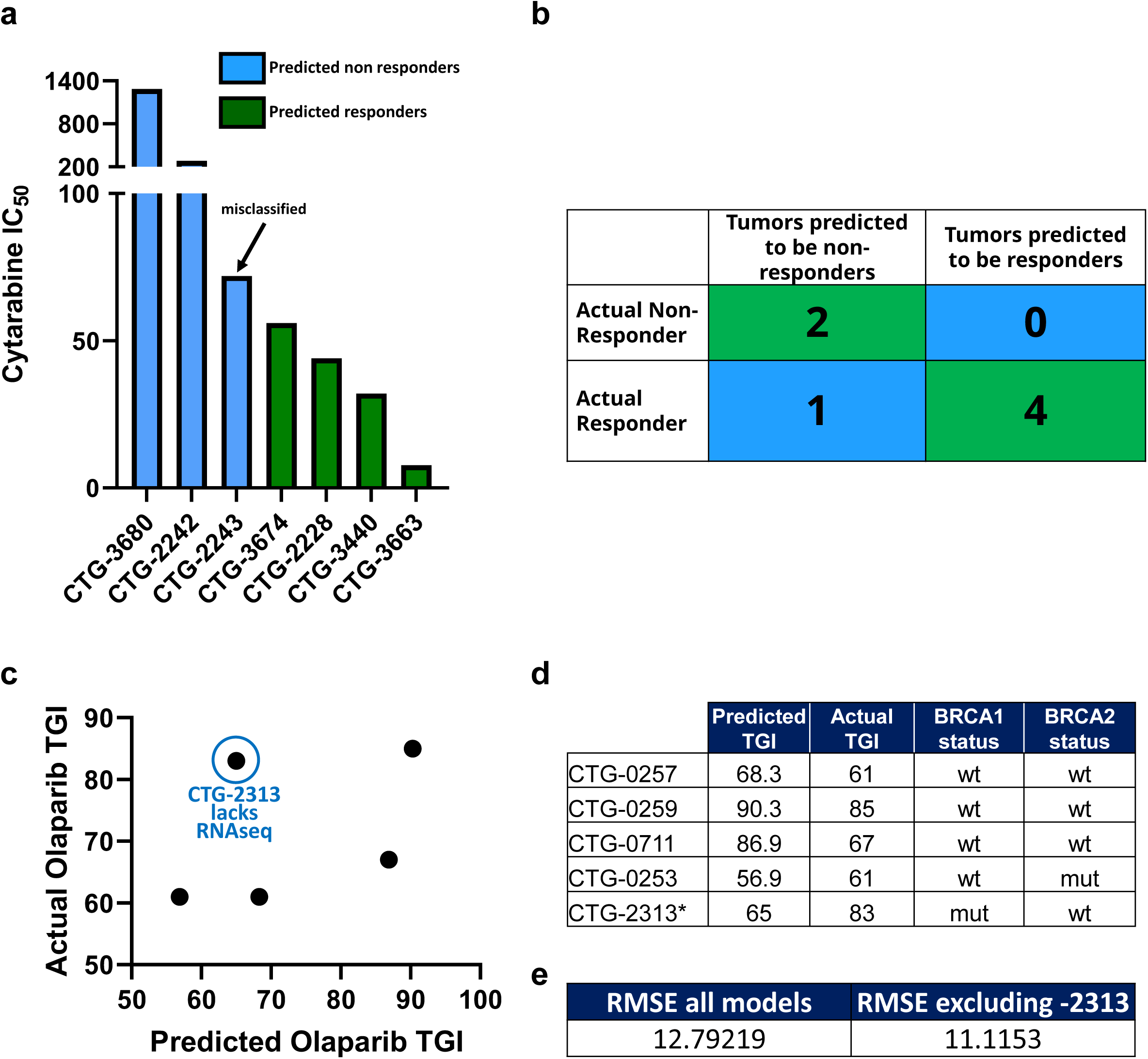
**(a)** Tumors are classified as Predicted NR (blue) and Predicted R (Green) based on their PPMO profile, and their actual IC_50_ values are plotted. **(b)** A confusion matrix of predicted vs actual R and NR tumors. **(c)** Predicted TGI values are plotted against TGI values obtained from *in vivo studies*. **(d)** Ovarian PDX models are shown with the predicted TGI, actual TGI and BRCA status. **(e)** Root-mean square error (RMSE) analysis of predicted vs actual TGI for all models in the cohort, and the cohort excluding CTG-2313.

The resulting Comp 1-Olaparib-based model was tested on 5 newly profiled Ovarian samples, which had molecular and response data; CTG-2313 lacked RNAseq data. Importantly, models with a diverse BRCA status were included in this exercise. Since this model was based on a regression analysis, we sought to predict TGI values to Olaparib therapy, rather than simple R vs NR categorization. Despite the somewhat low number of models used to generate the prediction, the computational model predicted TGI values with an RMSE of 12.8 when CTG-2313 was included, and 11.1 when CTG-2313 was excluded (Fig. 5c-e). Importantly, removing any omic block within the PPMO model led to a significant reduction in the predictive power, as noted by the higher error in TGI prediction for model CTG-2313 lacking RNAseq. Notably, prediction accuracy seemed completely independent of BRCA status (Fig.5d).

## Discussion

The two analyses presented here provide POC in the use of PPMO as the basis for next generation CDx assays that provide an accurate and prospective assessment of response profiles for a given therapeutic. The magnitude of these findings set the stage for using a more detailed approach in establishing CDx assays that are highly accurate and capture a broader patient population. Indeed, the dataset provided here is small in nature. However, given the success presented with this small cohort of tumors, we are confident that a scaling of the training set utilized would only strengthen the ability of PPMO to accurately provide prospective response predictions. Further, we acknowledge that the pharmacology input in this study was *in vivo-* and *ex vivo-*based readouts, rather than clinical outcomes. While significant research has demonstrated the clinical correlation between *in vivo* PDX modeling and clinical outcomes [25], evidence for an *ex vivo* correlation has not been substantiated. Nevertheless, as a POC, we view the exercise as one using a computational model to train on the complex molecule make up of a tumor based in a pharmacology variable input, to predict an identical pharmacology variable output. We speculate that changing the input/output variables to a clinical response would not impact the prediction accuracy. Further investigation in this matter will be revealing. Expansion to other tumor types and therapeutic modalities is also warranted.

Our integrative analysis revealed several interesting associations with tumors exhibiting sensitivity or resistance to cytarabine in AML. While both CD34+ and CD56+ associated with cytarabine response profiles, and had loadings on Comp 1-Cytarabine, only CD34+ populations showed strong correlations (*R >=* 0.8) with other molecular loadings of Comp 1-Cytarabine (Fig. 4a). Specifically, CD34+ populations levels positively correlated with the kinase activity of PDK1/2 and PDK3/4, which are known metabolic gatekeepers and negative regulators of oxidative respiration [26] (Fig. 2d inner, and outer lines, respectively). Importantly, while this profile was present in cytarabine sensitive tumors, it was absent in cytarabine resistant tumors (Fig. 2d). This suggests that tumors characterized by cytarabine resistance are enriched with a population of cells and exhibit increased oxidative respiration, while those tumors that are sensitive to cytarabine have an enrichment of CD34+ cell populations and exhibit a PDK-mediated inhibition of oxidative respiration. These results agree with a recent finding that high level of oxidative phosphorylation drives cytarabine resistance in AML [27]. In addition, a mutation in *RNF213* is top loading in Comp 1-Cytarabine (Fig. 2c) associating with cytarabine resistance, and recent studies have speculated that RNF213 serves as a metabolic gatekeeper. It was shown that RNF213 plays an important role in lipid metabolism and the modulation of lipotoxicity [28], as well as fat storage and lipid droplet formation [29]. Moreover, RNF213 associates with PTP1B and HIF1A to coordinate the cellular response to hypoxia and controls non-mitochondrial oxygen consumption [30]. We also found negative correlation between ALDH3B1 protein expression and PDK1/2 and PDK3/4 kinase activity, with ALDH3B1 protein expression enriched in NR tumors. Increased ALDH3B1 has been observed across tumor indications [31] and has been shown to increase ATP production [32]. Collectively, these results suggest a potential molecular switch occurring in CD34+ depleted tumors, potentially contributing to cytarabine resistance.

Our analysis found evidence for additional intriguing mechanisms that we speculate may be contributing to Cytarabine resistance in these tumors. Namely, we found a positive correlation between MYC copy number gain and SAMHD1 protein expression in NR tumors (Fig.3d). Indeed, recent studies have proposed SAMHD1 as a biomarker for Cytarabine resistance [18]. The proposed mechanism occurs via SAMHD1-mediated hydrolysis of Ara-CTP which depletes the cells of Cytarabine [18]. Additional reports have suggested that MYC acts to transcriptionally regulate SAMDH1 expression [17]. Taken collectively, these integrated finding support the MYC-SAMHD1 axis as a driver of and biomarker for cytarabine resistance. Further directed functional genomic investigations into the role of these biomarkers towards cytarabine resistance will be revealing.

Our integrative analysis also revealed several interesting associations with tumors exhibiting sensitivity or resistance to Olaparib in ovarian cancer. Interestingly, we found that reduced expression of LRIG1 protein expression associated with resistance to Olaparib (Fig.4c). This phenomenon has been observed in ovarian cancer cell lines exhibiting resistance to Etoposide [33]. We also found a downregulation of MEIS1 protein expression in tumors associating with reduced sensitivity to Olaparib (Fig.4c). MEIS1 has been implicated in the reorganization of epigenetic states via the recruitment of PARP1. Disruption of these epigenetic mechanisms, via the inhibition of PARP1, may render a tumor more sensitive than ones where these mechanisms are not active due to reduced MEIS1 expression. We also found increased protein expression of CIT and TAOK3 in tumors associated with reduced sensitivity to Olaparib (Fig.4c). CIT expression has been reported to provide protection against chromosome instability [34], while TAOK3 is involved in DNA repair via ATM-mediated p38 activation[35]. That increased protein expression of the phosphate efflux transporter, XPR1, is associated with Olaparib response was intriguing (Fig.4c). A recent study has shown a vulnerability in ovarian cancer cell lines when XPR1 activity is inhibited when Solute Carrier Family 34 Member 2 (SLC34A2) is overexpressed do to intracellular phosphate overload [20].

Collectively, these findings support the power of using PPMO to better understand tumor cell biology and its relationship to pharmacology profiles. By leveraging a PPMO in this study, we identified a biomarker profile which may contain targets suitable for therapeutic intervention. Further analyses into the veracity of these biomarkers as therapeutic targets will be revealing. The biomarkers identified in this comprehensive study are intriguing and warrant further evaluation into potential roles they play in conferring therapeutic resistance. Importantly, none of the biomarkers presented can predict response profiles independently of the entire biomarker profile. Accordingly, our findings demonstrate an early proof on concept on the strength of using PPMO models to establish diagnostic or companion diagnostic platforms.

## Methods

### Human acute myeloid leukemia specimens

AML patient specimens were obtained from the Stem Cell and Xenograft Core at the Perelman School of Medicine, University of Pennsylvania. The consent to collect and use human AML specimens for research was obtained under the protocol “Hematologic Malignancies Tissue Bank”, which is approved by the University of Pennsylvania’s Institutional Review Board (IRB protocol #703185).

### Human leukapheresis sample processing and cryopreservation

Leukapheresis from AML patients were collected at the Perelman Center for Advanced Medicine, Cancer Center, Hospital of University of Pennsylvania, in pheresis bag and transported to the lab for further processing within 2 hours of sample collection. Briefly, cells were transferred in 50ml Falcon tubes (Fisher), counted using a Nexcelom Cell Counter (Nexcelom) and diluted with PBS 2% fetal bovine serum (Gemini Bio) to reach a concentration of 100 to 200 million cells/ml. After centrifugation, red blood cells were lysed using ammonium chloride (Stem Cell Technologies) and cells were washed and resuspended in PBS 2% FBS for counting. Cells were then aliquoted and frozen at an appropriate cell concentration in a mix 1:1 of PBS 2% FBS and 14% dimethyl sulfoxide (Fisher), 4% Hetastarch (NovaPlus), and 4% bovine serum albumin (Gemini Bio). Cell vials (Abdos) were placed at -80C overnight and transferred into liquid nitrogen freezer for long-term conservation.

### Human cells

All patient samples were procured after informed consent and deidentified in accordance with IRB-approved protocols. White blood cells were obtained via leukapheresis from AML patients in blast crisis. Mononuclear cells were separated by density gradient centrifugation, washed and then cryopreserved. Cells were stored in vapor phase liquid nitrogen until use for ex vivo assay (flow cytometry, cytotoxicity assays and molecular profiling).

### Flow Cytometry Method

One comprehensive 18-color panel of monoclonal antibodies was used to stain cells for AML characterization. The following surface antibodies were used in the panel (Supplementary Table 2): anti-CD14 BUV395 (clone M5E2), anti-CD4 BUV496 (clone RPA-T4), anti-CD34 BUV661 (clone 581), anti-CD19 BUV737 (clone SJ25C1), anti-CD123 BV421 (clone 9F5), anti-CD3 BV605 (clone SK7), anti-CD38 BV650 (clone HIT2), anti-CD64 BV711 (clone 10.1), anti-CD13 BB515 (clone WM15), anti-CD11b BB700 (clone ICRF44), anti-CD7 PE (clone M-T701), anti-CD15 AF647 (clone W6D3), all from BD Bioscience, and anti-CD45 BV510 (clone HI30), anti-HLA-DR BV570 (clone L243), anti-CD56 PE-Dazzle594 (clone 5.1H11), anti-CD33 PE-Cy7 (clone P67.7) and anti-CD117 APC-Cy7 (clone 104D2), all from Biolegend. In addition, a fixable viability dye in the FVS700 channel was used to exclude non-viable cells in the gating strategy.

Prior to staining the patient samples, each antibody was titrated using an 8-point serial dilution beginning with twice the manufacturer’s recommended concentration. Separation index was calculated, and the optimal antibody amounts were determined. Fluorescence Minus One (FMO) controls and Fluorescence Minus X (FMX) controls were also prepared and analyzed during panel optimization to establish the optimal gating strategy and to set gates properly. 0.5 × 10^6^ thawed, washed cells were plated per well and viability dye was added and incubated for 15 (± 5) minutes. Cells were washed twice in FACS stain buffer then blocked for 10-15 minutes with Fc block. Post blocking, conjugated antibodies were added and incubated, protected from light for 30-40 minutes, then washed in FACS stain buffer twice before resuspending in FluoroFix buffer (Biolegend). Cells were incubated for 20 minutes protected from light, and then washed once more in FACS stain buffer. Cells were resuspended in FACS stain buffer and stored at 2-8°C, protected from light, until acquisition on the flow cytometer.

### Instrument Calibration and Compensation

A 5-laser, 26-parameter, Becton Dickinson FACS Symphony A3 (4-Blue, 3-Red, 7-Violet, 4-Gr, 6-UV, see Table x for cytometer configuration) with BD FACSDiva vxx software was used for sample acquisition. Prior to sample acquisition, the instrument was calibrated and fluidics were quality controlled using BD CS&T beads. Once it was confirmed that CS&T beads passed acceptance, compensation was performed to adjust for spectral overlap within each channel. Single-color stained beads were acquired on the cytometer, and compensation was calculated using the instrument software, and then applied to the samples.

### Flow Cytometry Data Collection and Analysis

Cytometer configuration is shown in supplementary table 3. 500,00 viable cells were collected from each sample. FCS files were exported and analyzed with BD Biosciences FlowJo v10 software. Cells were first gated from SSC-A versus FSC-A to exclude debris, then gated from FSC-H versus FSC-A, then SSC-H versus SSC-A to include only singlets. Then cells were gated from SSC-A versus Viability FVS700, and negative cells gated to include only viable cells and to exclude all dead cells. At this point, the gating strategy in supplementary table 4 was used to identify CD45 blast cells, AML blast cells, AML blast progenitor cells, AML blast leukemic stem cells (LSC), AML monoblasts, leukocytes, NK cells, T cells, Helper T cells, Cytotoxic T cells, and B cells.

### Cytotoxicity assay

Primary patient leukapheresis specimens were thawed, washed twice in media, and counted using acridine orange and propidium iodide with Cellometer Auto2000 (Nexcelom, Lawrence, MA). Cells were seeded in 96-well plates in Champion’s proprietary AML VitroScreen media. Cytarabine, prepared in DMSO stock solution, was serially diluted in AML VitroScreen media, or media alone and then added to wells in triplicates. Cells were incubated for 6 days in a 37°C 5% CO2 humidified incubator. On day 6, Cell Titer Glo reagent (Promega) was equilibrated to room temperature, then added to each well and incubated for 10 minutes with shaking before plate reading. Luminescence was recorded using Infinite M Plex plate reader (Tecan). Cell viability IC_50_ curves were generated from log transformed, normalized luminescent (RLU) readings with nonlinear regression (4PL curve fit) using Graph Pad Prism 8 to establish IC_50_, and R^2^ values.

### *In vivo* pharmacology studies

All methods were carried out in accordance with relevant guidelines and regulations for using animals in the study. All studies involving animals were reviewed and approved by the Institutional Animal Care and Use Committee at Champions Oncology, Inc. Tumor volumes are recorded for each experiment beginning seven to ten days after implantation into Nude mice. When tumors reach an average tumor volume of 150-300mm^3^ animals are matched by tumor volume into treatment or control groups to be used for dosing and dosing initiated on Day 0. Animals are visually examined daily. Tumor volumes are taken twice weekly. A final tumor volume is taken on the day study reaches endpoint. Animals are weighed twice weekly. Animals are weighed twice weekly. A final weight is taken on the day the study reaches end point or if animal is found moribund, if possible. The study endpoint is when the mean tumor volume of the control group (uncensored) reaches 1500mm^3^. If this occurs before Day 28, treatment groups and individual mice are dosed and measured up to Day 28. TGI is calculated as 1 – (mean volume of treated tumors)/(mean volume of control tumors)) × 100%. Olaparib was dosed orally at 100 mg/kg once a day for 28 days.

### Molecular Profiling

#### Stranded mRNA Library Preparation

Qualified RNA was used for library preparation using Illumina Stranded mRNA Prep ligation. first, mRNA was purified and captured with Oligo(dT) magnetic beads. Second, purified mRNA was fragmented and copied into first strand complimentary DNA (cDNA) using reverse transcriptase and random primers, in the following steps, dUTP replaced dTTP to form strand specific cDNA. In the final steps adenine(A) and thymine (T) bases were incorporated into fragment end and ligate adapters. The resulting libraries were purified and selectively amplified for sequencing.

#### Gene Quantification pipeline

Rna-Seq raw reads were preprocessed with adapters and low quality bases trimming using Cutadapt[36], following by alignment to Hg19 human genome and GRCh37.p19 as a gene model by STAR aligner[37]. Genes counts were extracted by RSEM[38] followed by TMM depth normalization[39] and removal of known batch effects via ComBat[40]. Low coverage genes were removed and log TPM scores were calculated for every gene.

#### Exome library preparation

Extracted DNA was fragmented using provided enzyme in the kit (Agilent SureSelect XT HS2 DNA system), the fragmented DNA then went through End repair and dA-tailing followed by adaptor ligation to add molecular barcode. The barcoded library was purified, amplified, and followed by quality checks to make sure enough libraries were generated for hybridization using SureSelect Human All Exon V7 probe. The hybridized libraries were then captured using streptavidine-coated beads and amplified for sequencing.

#### Low Coverage Variants Identification

To augment WES variant calling an amplicon approach was also applied on 54 genes using TruSight myeloid sequencing panel[41] (Illumina, CA) and paired-end sequencing runs performed on a MiSeq (Illumina) genome sequencer. Sequences obtained were analyzed by GATK best practices and annotated by Ensembl Variant Effect Predictor[42].

#### WES Variant Calling and Copy Number Identification

A comprehensive variant calling pipeline was used to overcome the lack of a match normal sample. Briefly, multiple callers were used for SNPs (Mutect, LoFreq and snp-strelka) and INDELs (indel-Strelka, Pindel and Scalpel) with a conjugated diploid cell line NA12878 as a reference sample. High quality variants with an agreement of two callers were qualified and known germlines with population prevalence’s in ExAc database[43] above 1% were excluded. Copy number were identified via Excavator[44] pipeline using NA12878 as the normal reference.

#### Multi Omics Clustering

RNA-seq genes, proteomics and phosphor-proteomics blocks were used to cluster AML samples. For every block 2000 of the most variable genes/proteins were selected and an integrative clustering analysis was applied using Bayesian latent variable identification via iClusterPlus[45] using gaussian priors, 180k burn in and 300K draws and k=2-11. Hierarchal clustering of Latent variables and average silhouette used to identify k=3 as the most robust number of clusters.

### Proteomics and Phosphoproteomics Profiling

#### Sample Lysis

Total 47 were received. About 30M AML cells were mixed with 500 μL of lysis buffer that is 9M urea, pH 8.5 20mM HEPEs. Samples were water bath sonicated for 30mins follow by spinning in the high-speed centrifuge for 10mins at 14,000 rpm. To complete the lysis, samples were supersonicated for 30 seconds at 20% amplitude (Qsonica, Q500 Sonicator) Spin down samples with tabletop centrifuge. Protein concentration of samples were measured by BCA assay (Cat No: A53225, ThermoFisher Scientific) post sample lysis.

#### Proteomics Sample Preparation

About 2 mg of each sample was taken from the lysate and normalize to the same volume with lysis buffer. Samples were reduced in 10 mM DTT for 25 min at 60 °C, and then reduced samples were alkylated in 20 mM IAM in dark environment for 20 mins at room temperature. Excess IAM in the samples were quenched by adding 100 mM DTT solutions. DI water and HEPE pH 8.5 were added to each sample so that final urea concentration was diluted to 1.6 M, and final pH about 8 for enzymatic digestion. 40 μg of Try/LysC (Cat No: A41007, ThermoFisher Scientific) were added to each sample. Samples were incubated overnight at 37 °C for 12 hours. Additional 10 μg of Tryp/LysC was added to each sample the next day and samples were incubated for 4 more hours to complete the enzymatic digestion.

#### Peptide Cleanup

10% TFA were added into each digested peptides samples and form a final concentration of 1% TFA. pH was tested and acidic. Then, acidified samples went through 100 mg SEK PAK column (Cat No: 60108-302, ThermoFisher Scientific) for desalting. 25% of desalted peptides aliquot (0.5mg of equivalent protein content) of each sample was taken for normal DIA analysis, and the remaining 75% of desalted peptides of each sample (1.5mg of equivalent protein content) was reserved for phospho-DIA analysis. Within normal DIA sample, 80% of each sample (0.4mg of equivalent protein content) were taken to pool together for peptide fractionation; remaining 20% of each sample was dried and stored for LC/MS analysis.

#### Phospho-peptides enrichment

Reserved peptides were enriched with High-Select™ TiO_2_ phosphopeptides enrichment kit from Thermo Scientific (Cat No: A32993, ThermoFisher Scientific) using protocol from Thermo Scientific with the exception for the elution buffer (80% ACN, 5% Ammonium Hydroxyl). Within phospho-DIA enriched samples, 80% of each sample (1.2mg of equivalent protein content) were taken to pool together for peptide fractionation; remaining 20% of each sample (0.3mg of equivalent protein content) was dried and stored for LC/MS analysis.

#### Peptide Fractionation

Both the normal DIA and phospho-DIA library composite samples were fractionated into 96 fractions with a high pH reverse phase offline HPLC fractionator (VanquishTM, ThermoFisher Scientific). Mobile phase A is DI H_2_ O with 20 mM Formic Acetate, pH 9.3; mobile phase B is Acetonitrile (Optima™, LC/MS grade, Fisher Chemical™) with 20mM Formic Acetate, pH 9.3. Gradient of separation is displayed in Supplementary Table 6. 96 fractions were then combined into 24 fractions and ready for Liquid Chromatography Mass Spectrometry (LC/MS) analysis.

#### LC/MS Analysis

All fractionated samples were analyzed by nano flow HPLC (Ultimate 3000, Thermo Fisher Scientific) followed by Thermo Orbitrap Mass Spectrometer (QE HF-X). Nanospray Flex™ Ion Source (Thermo Fisher Scientific) was equipped with Column Oven (PRSO-V2, Sonation) to heat up the nano column (PicoFrit, 100 μm x 250 mm x 15 μm tip, New Objective) for peptide separation. The nano LC method is water acetonitrile based 150 minutes long with 0.25 μL/min flowrate. For each library fractions, all peptides were first engaged on a trap column (Cat. No: 160454, Thermo Fisher) and then were delivered to the separation nano column by the mobile phase. A specific of gradient information was indicated in Supplementary Table 7. For DDA library construction, a DIA library specific DDA MS2-based mass spectrometry method on Eclipse was used to sequence fractionated peptides that were eluted from the nano column. For the full MS, 120,000 resolution was used with the scan range of 375 m/z – 1500 m/z. For the dd-MS(MS2), 15,000 resolution was used, and Isolation window is 1.6 Da. ‘Standard’ AGC target and ‘Auto’ Max Ion Injection Time (Max IT) were selected for both MS1 and MS2 acquisition. Collision Energy (NCE) was set to 35%, and total cycle time is 1 sec. For DIA analytical samples, a high-resolution full MS scan followed by two segment DIA methods was used for the DIA data acquisition. For the full MS scan, 120,000 resolution was used for the range of 400 m/z – 1200 m/z with ‘Standard’ AGC target and 50 ms Max IT. For both DIA segments, details of isolation windows (IW) and precursor mass range are shown in Supplementary Table 8 & Supplementary Table 9. For DIA fragments scan, 30,000 resolution was used for the range of 110 m/z – 1,800 m/z with ‘Standard’ AGC target and ‘Auto’ Max IT.

### Bioinformatic Analysis Pipeline Overview

This process is based on the sample data generated from a high-resolution mass spectrometer. DDA data was identified by Andromeda search engine within MaxQuant, and Spectronaut™ used identification results for spectral library construction. MaxQuant was used for identification of DDA data, served as a spectrum library for subsequent DIA analysis. MaxQuant was also used to localized Phosphosites for phosphopeptides. The analysis used raw data as input files and set corresponding parameters and human databases (UP000005640), then performed identification and quantitative analysis. The identified peptides satisfied FDR <=1% will be used to construct the final spectral library. For this DIA dataset, Spectronaut™ was used to construct spectral library information to complete deconvolution and extraction, and then mProphet algorithm was used to complete analytical quality control (1% FDR) to obtain reliable quantitative results. GO, COG, Pathway functional annotation analysis and time series analysis were also performed in above pipeline. DIA and phospho-DIA quantification data was first processed by dataProcess function in MSstats package, where equalize medians normalization and Tukey’s median polish summarization were used as default. MStats, which core algorithm is linear mixed effect model, processed DIA and phospho-DIA quantification result data according to the predefined comparison group, and then performed the significance test based on the model. Thereafter, differential protein and phospho-proteins screening was performed, and fold change ≥ 2 and adj p-value < 0.05 was defined as significant difference. Based on the quantitative comparison results, the differential proteins and phospho-proteins between comparison groups were found, and finally function enrichment analysis, protein-protein interaction (PPI) and subcellular localization analysis of the differential proteins were performed.

Proteomics data were normalized by variance stabilizing transformation, and missing values were imputed using a bayesian (‘bpca’) or a left-censored random draw (‘MinProb’) method by the R ‘DEP’ package[46].

### Kinase activity calculation

Kinase activity was calculated from the phosphoproteomics data by the ssGSEA algorithm [<u>19847166</u>]. Known phosphorylation links were compiled from 3 databases (“Signor” [<u>31665520</u>], “PhosphositePlus” [<u>31345222</u>] and “PDTs” [https://doi.org/10.1038/s41587-019-0391-9]), and were used as gene sets. The consolidated list consisted of 435 kinases and links to 11,022 phosphorylation substrates. The enrichment analysis was applied over our phospho-proteomics data using the consolidated list and kinases with identical targets in the data were collapsed, yielding 184 kinase entities. Enrichment scores were mean centered and normalized within each set across the samples.

### Multi Omics analyses

The input data consisted of 6 blocks: RNA expression, proteomics, kinase activity, mutated genes, genes with copy-number variations (CNVs) and cell sorting data. Mutated genes and genes with CNVs were included in downstream analysis only if they were present in the Cancer Gene Census (COSMIC) [https://cancer.sanger.ac.uk/census]. Features from numeric blocks (RNA, proteomics and kinase activity) were filtered by variance, with the threshold: min(“median peak”, median + MAD(gene variance)), where “median peak” is defined as 2*median (gene variance) - min(gene variance). The feature counts before and after filtering are shown in figure 1. Multi-omics data factorization and modelling was performed by sparse partial least squares discriminant analysis (sPLS-DA), implemented in the “mixOmics” R package [<u>29099853</u>]. To select features from each block, which optimize discrimination between cytarabine responders (IC50 < 100) and non-responders (IC50 > 100), a tuning step was run over series of counts, where each iteration attempts to optimize the discrimination performance using a number of features equal to the input counts. Thus, for each block, discrimination performance was measured on 15, n/10 and n/7 features. For RNA and proteomics 30 and 70 features were also included. The parameters used for classification error estimation during training were Mahalanobis distance as a distance metric, balanced error rate (BER), Horst optimization scheme, and leave-one-out (loo) validation. The inter-block association matrix was set to “null for discrimination purposes, and to “full” for feature correlation and network reconstruction. The optimal number of features per block per component are shown in table 1.

### Gene network

Mutual information (MI) matrix was calculated over values of features selected by sPLS-DA, along with cytarabine IC50 values by parallel estimation of the MI of vectors using entropy estimates from K-nearest neighbor distances [https://pubmed.ncbi.nlm.nih.gov/15244698/]. The gene network was then calculated by Context Likelihood or Relatedness (CLR algorithm, https://pubmed.ncbi.nlm.nih.gov/17214507/).

## Figures

**Supplementary Figure 1:**
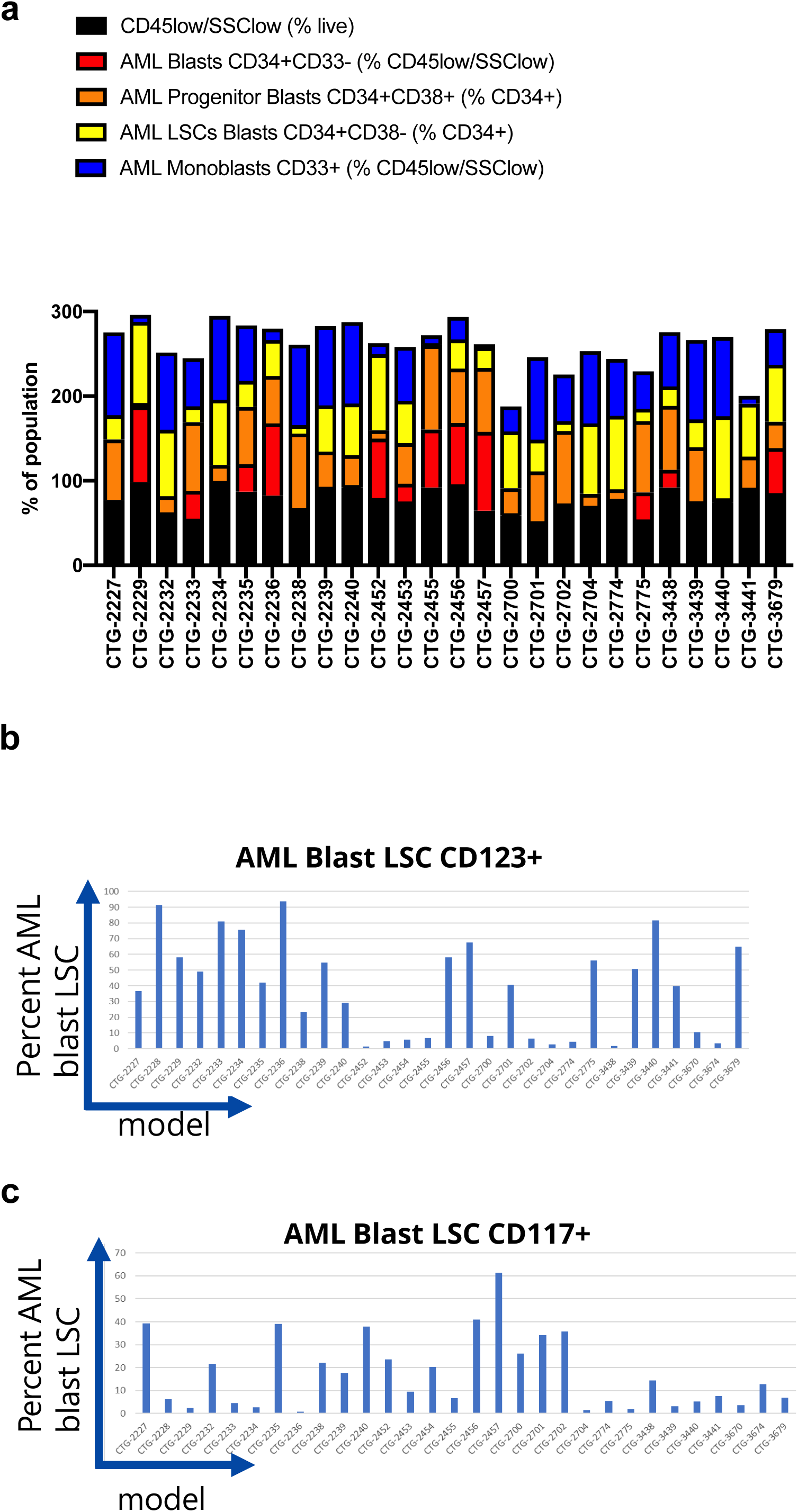
**a**. the summary of PhenoSeek analysis for AML Blasts, Progenitor Blast, LSC Blasts and Monoblasts are show for each sample are shown. The distribution of CD123^+^ **(b)** and CD117^+^ **(c)** LSCs across all samples are shown.

**Supplementary Figure 2:**
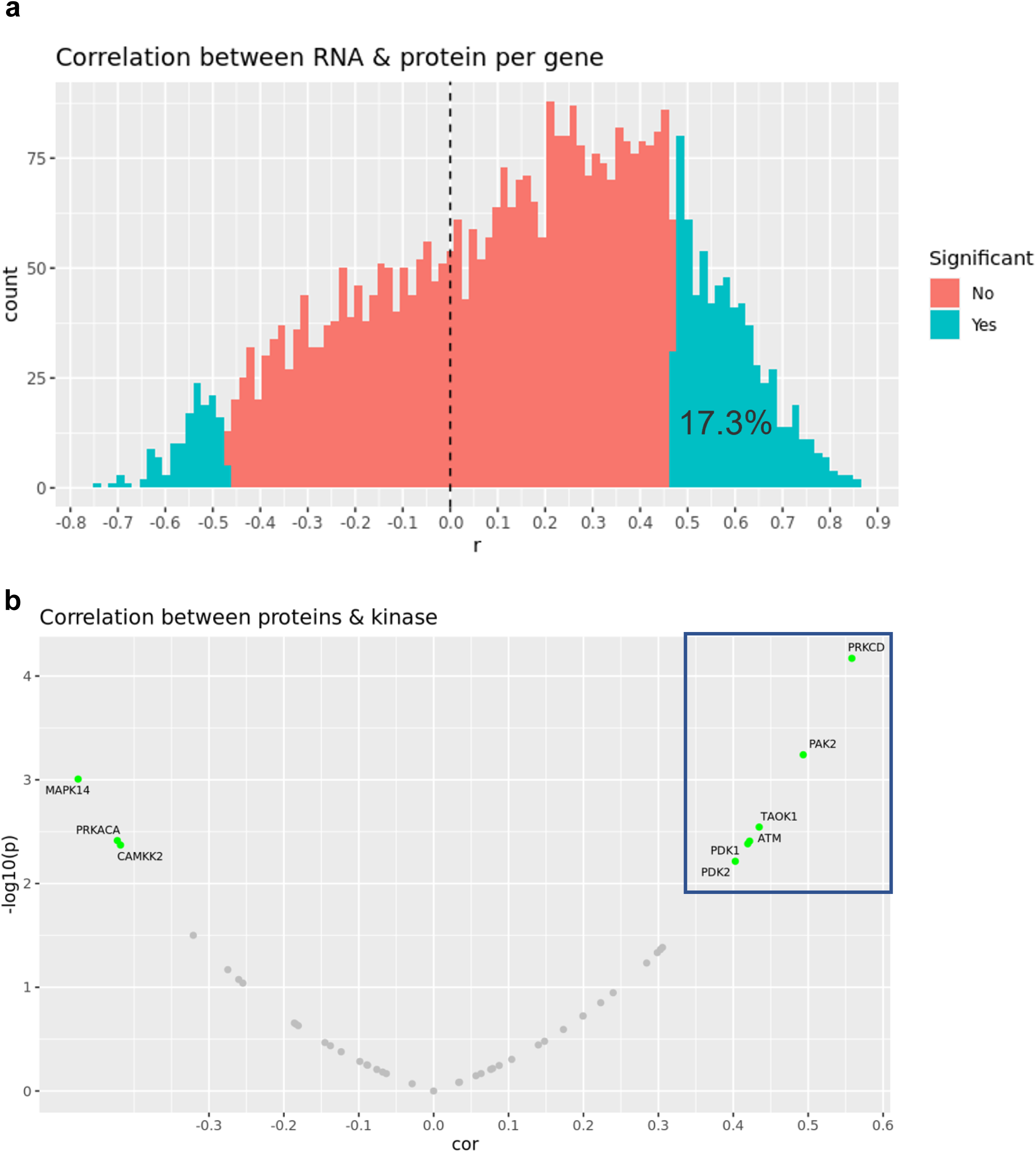
**a**. Correlations between proteins and their transcripts are positive and significant only for 17.3% of genes. **b**. Correlations between kinase activities and their expression levels. Positive correlations suggest the activity is driven by the protein expression.

**Supplementary Figure. 3:**
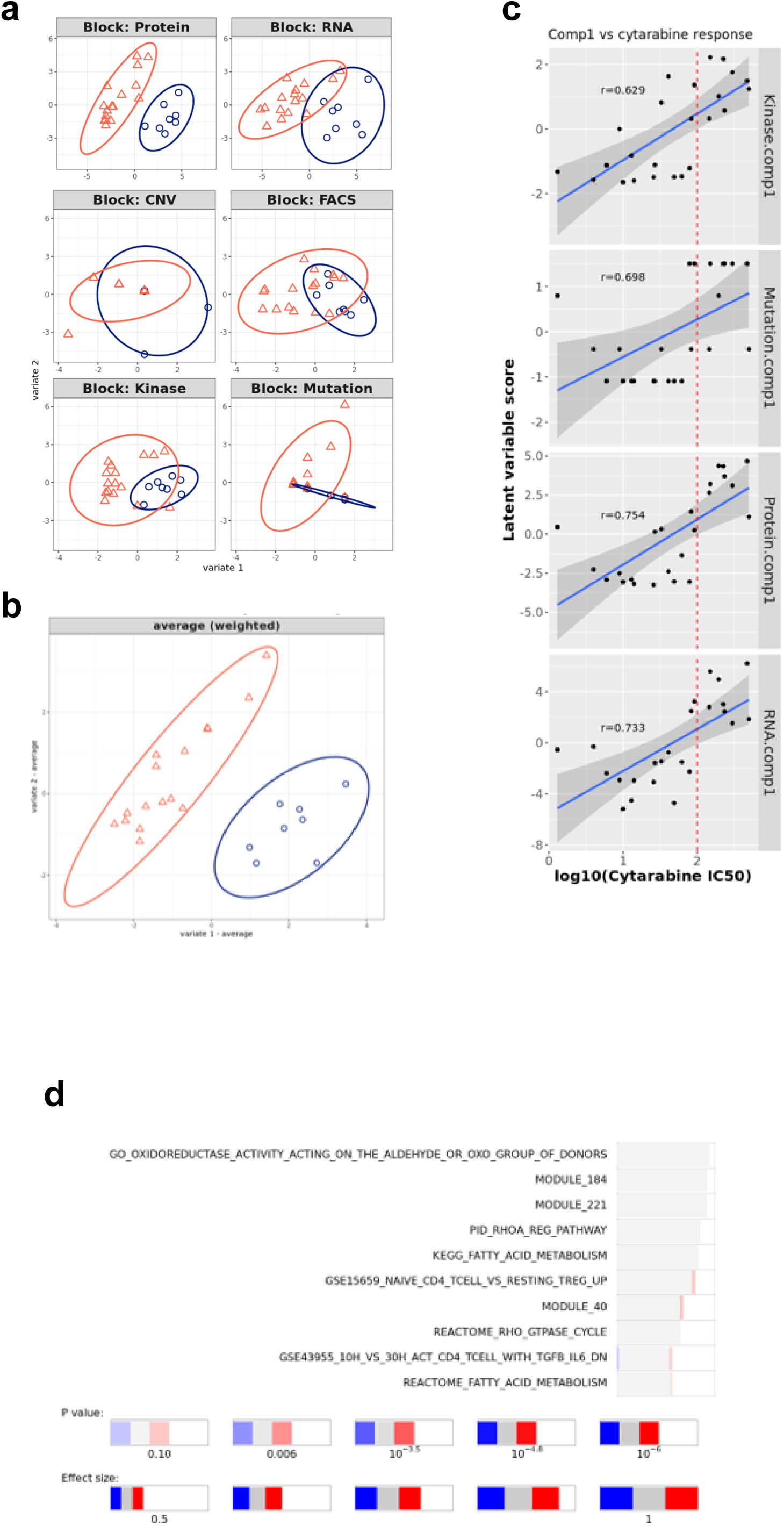
**a**. Plots of latent variables 1,2 for each omics block. Responders are shown in orange triangles, and non-responders in blue circles. **b**. Plots for weighted means of latent variables 1,2 from all omics blocks. **c**. Scatter plots and correlations between the first component of the different blocks and cytarabine IC50. The vertical red line indicates the response criterion. **d**. Functional enrichment analysis calculated for proteins and transcripts ranked by their absolute Comp1 loadings. The effect size is measured in area under curve (AUC). Terms with AUC > 0.65 and adjusted-p < 0.1 are shown. Blue: proportion of genes with negative loadings. Red: proportion of genes with positive loadings.

**Supplementary Figure. 4:**
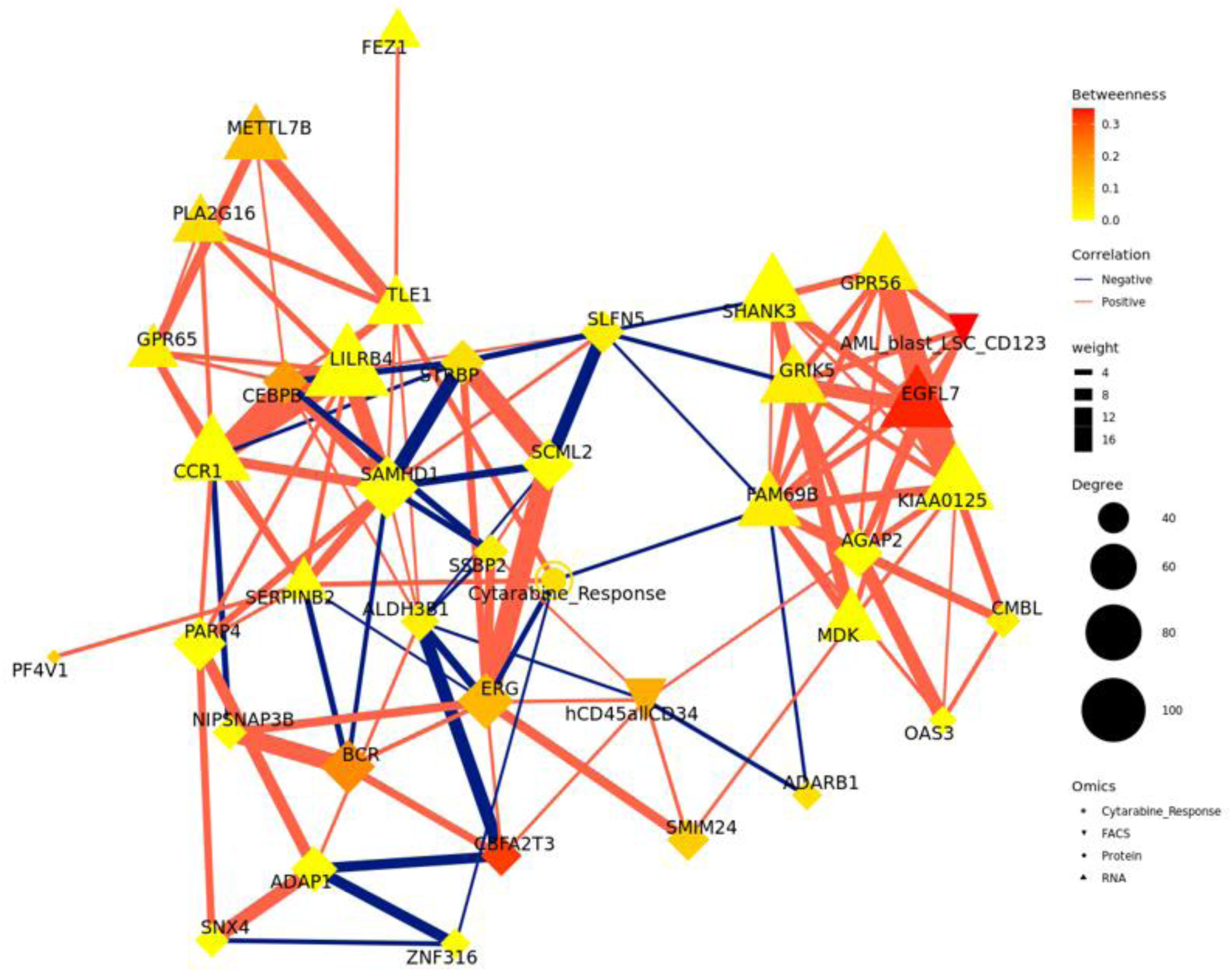
Context Likelihood or Relatedness network (CLR) calculated over multi-omics features from component 1, along with cytarabine response IC_50_. Shown here is a 2-neighborhood subgraph around cytarabine response, filtered for strongest (edge weight > 2) and significant (correlation FDR < 0.05) relationships. Edge thickness indicates interaction strength (weight).

**Supplementary Figure 5:**
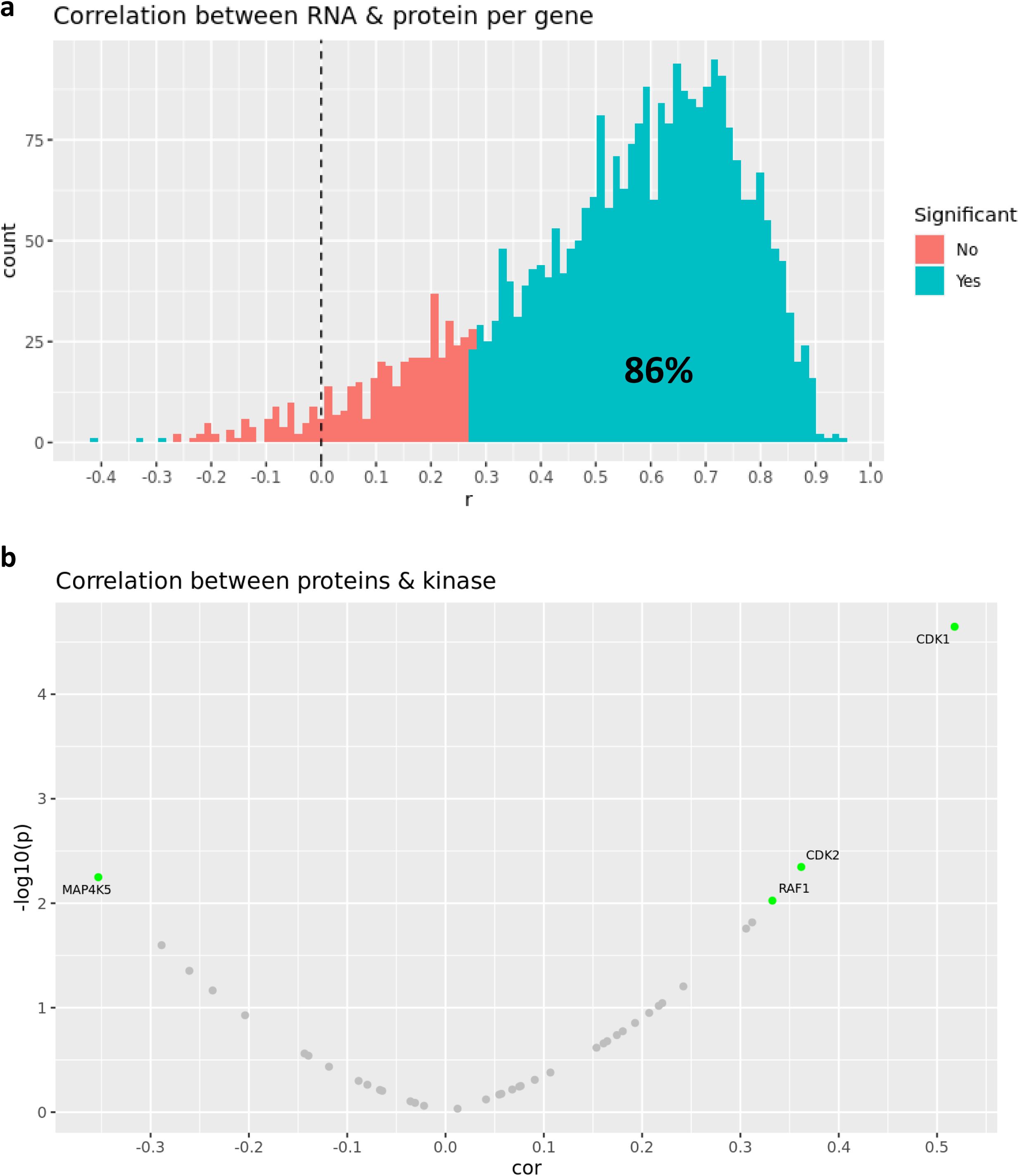
**(a)** Correlations between proteins and their transcripts are positive and significant for 86% of genes. (b) Correlations between kinase activities and their expression levels. Positive correlations suggest the activity is driven by the protein expression.

**Supplementary Fig 6.**
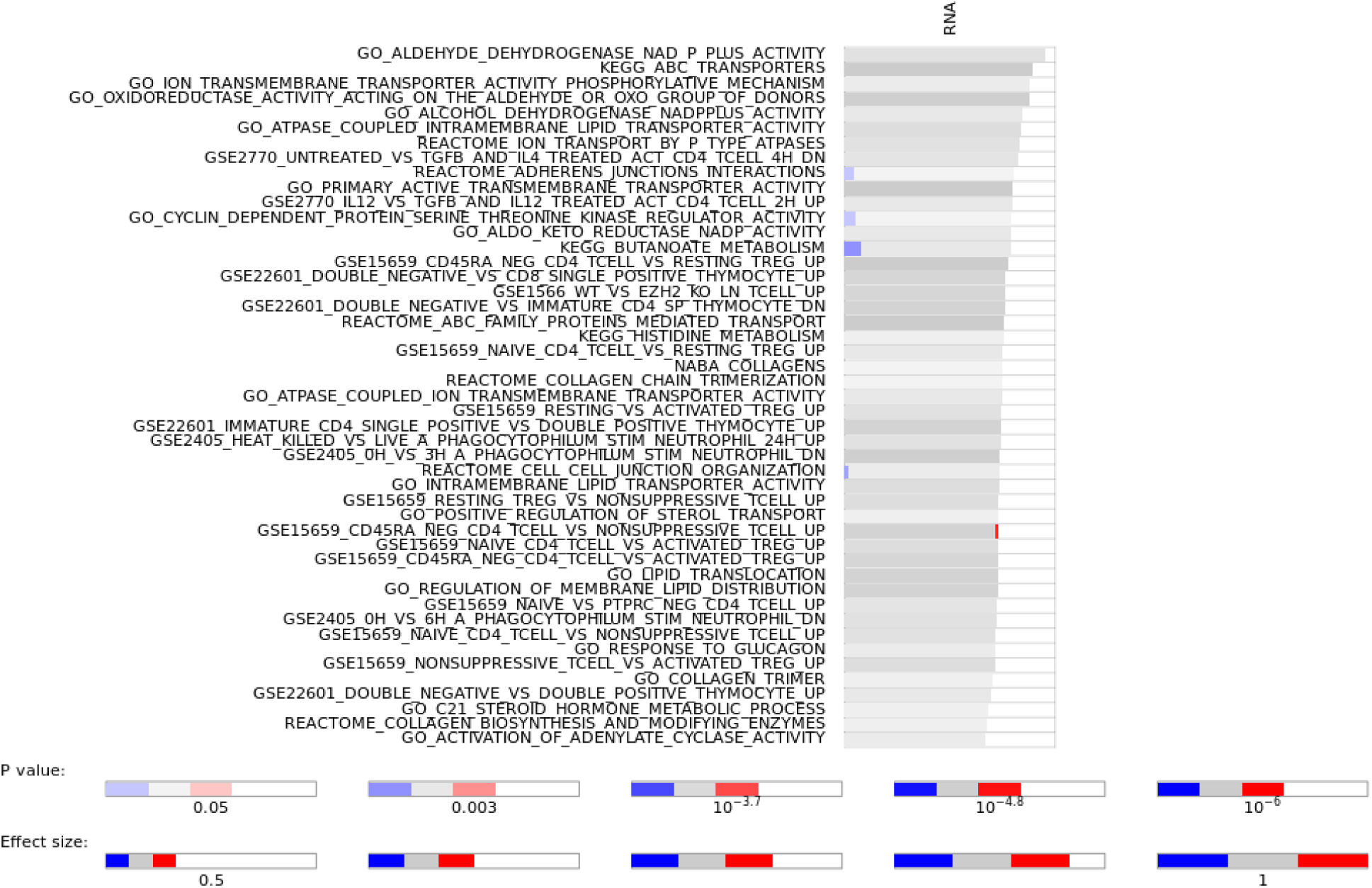
Functional enrichment analysis calculated for proteins and transcripts ranked by their absolute Comp1 loadings. The effect size is measured in area under curve (AUC). Terms with AUC > 0.65 and adjusted-p < 0.1 are shown. Blue: proportion of genes with negative loadings. Red: proportion of genes with positive loadings.

---

**Supplementary Table 2:** *Phenoseek* Gating Strategy.

| Cell Subset | Parent | Immunophenotype |
| --- | --- | --- |
| CD45 blast | Viable | SSC<sub>low</sub>, CD45<sub>low</sub> |
| AML blast | CD45 blast | CD34+CD33- |
| AML blast progenitor | AML blast | CD34+CD38+ |
| AML blast progenitor markers | AML blast progenitor | CD123+ |
| AML blast LSC | AML blast | CD34+CD38- |
| AML blast, LSC markers | AML blast LSC | CD117+, CD123+ |
| CD45 blast markers | CD45 blast | CD4+, CD7+, CD11b+, CD13+,<br>CD14+, CD15+, CD56+, CD64+,<br>HLA-DR+, CD19+ |
| AML monoblast | CD45 blast | CD33+ |
| AML monoblast markers | AML monoblast | CD4+, CD7+, CD11b+, CD13+,<br>CD14+, CD15+, CD56+, CD64+,<br>HLA-DR+, CD19+ |
| Leukocytes | Viable | CD45+/<sub>hi</sub> |
| NK cells | CD45+/<sub>hi</sub> | CD3-CD56+ |
| T cells | CD45+/<sub>hi</sub> | CD3+CD19- |
| Helper T cells | CD3+CD19- | CD3+CD4+ |
| Cytotoxic T cells | CD3+CD19- | CD3+CD4- |
| B cells | CD45+/<sub>hi</sub> | CD3-CD19+ |

**Supplementary Table 3:**
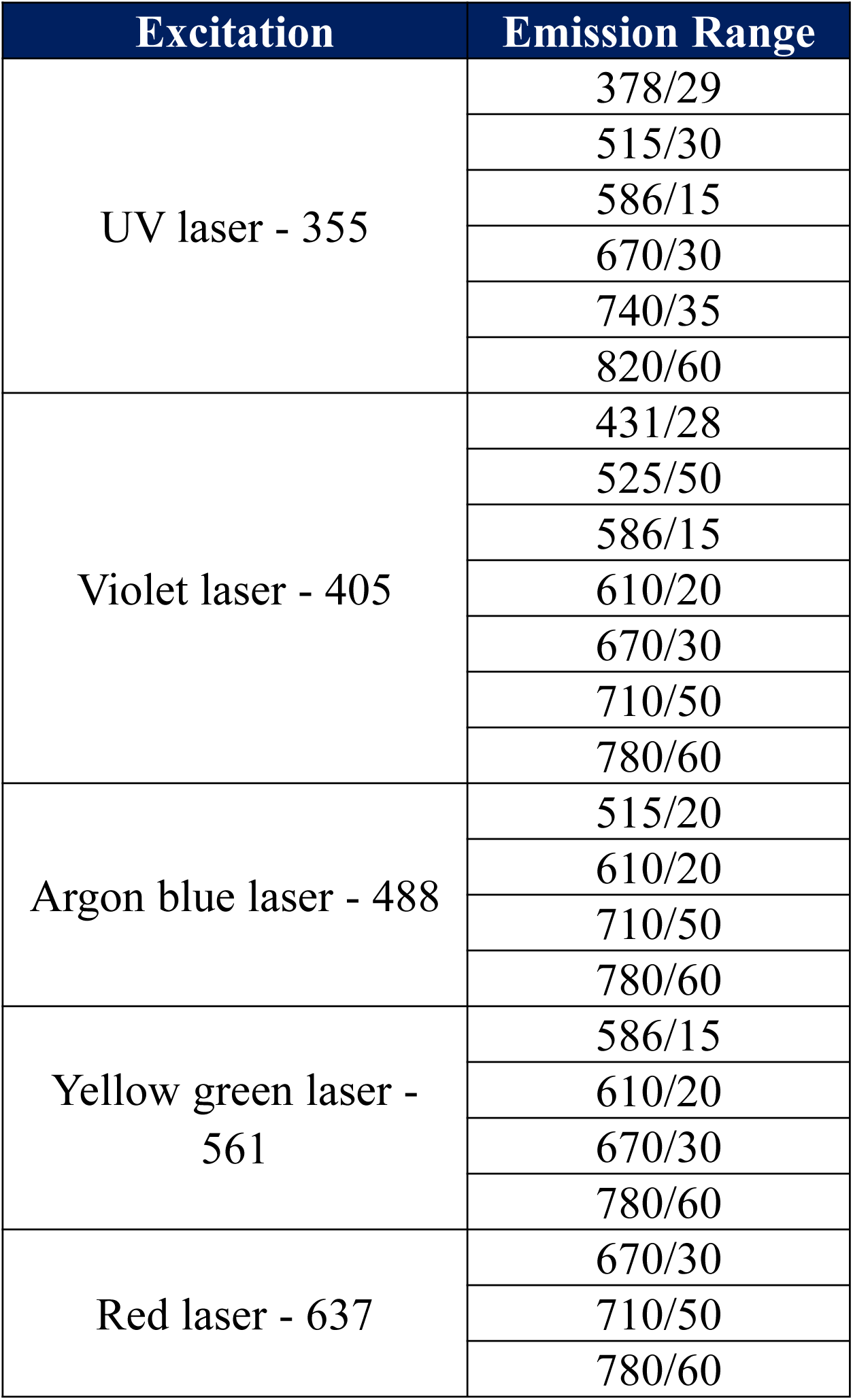
*Phenoseek* Cytometer Configuration.

**Supplementary Table 4:** Panel List.

| Marker | Fluorophore | Clone | Supplier |
| --- | --- | --- | --- |
| CD14 | BUV395 | M5E2 | BD Biosciences |
| CD4 | BUV496 | RPA-T4 | BD Biosciences |
| CD34 | BUV661 | 581 | BD Biosciences |
| CD19 | BUV737 | SJ25C1 | BD Biosciences |
| CD123 | BV421 | 9F5 | BD Biosciences |
| CD45 | BV510 | HI30 | BioLegend |
| HLA-DR | BV570 | L243 | BioLegend |
| CD3 | BV605 | SK7 | BD Biosciences |
| CD38 | BV650 | HIT2 | BD Biosciences |
| CD64 | BV711 | 10.1 | BD Biosciences |
| CD13 | BB515 | WM15 | BD Biosciences |
| CD11b | BB700 | ICRF44 | BD Biosciences |
| CD7 | PE | M-T701 | BD Biosciences |
| CD56 | PE-Dazzle594 | 5.1H11 | BioLegend |
| CD33 | PE-Cy7 | P67.6 | BioLegend |
| CD15 (SSEA-1) | AF647 | W6D3 | BioLegend |
| Viability | FVS700 | NA | BD Biosciences |
| CD117 | APC-Cy7 | 104D2 | BioLegend |

**Supplementary Table 6.**
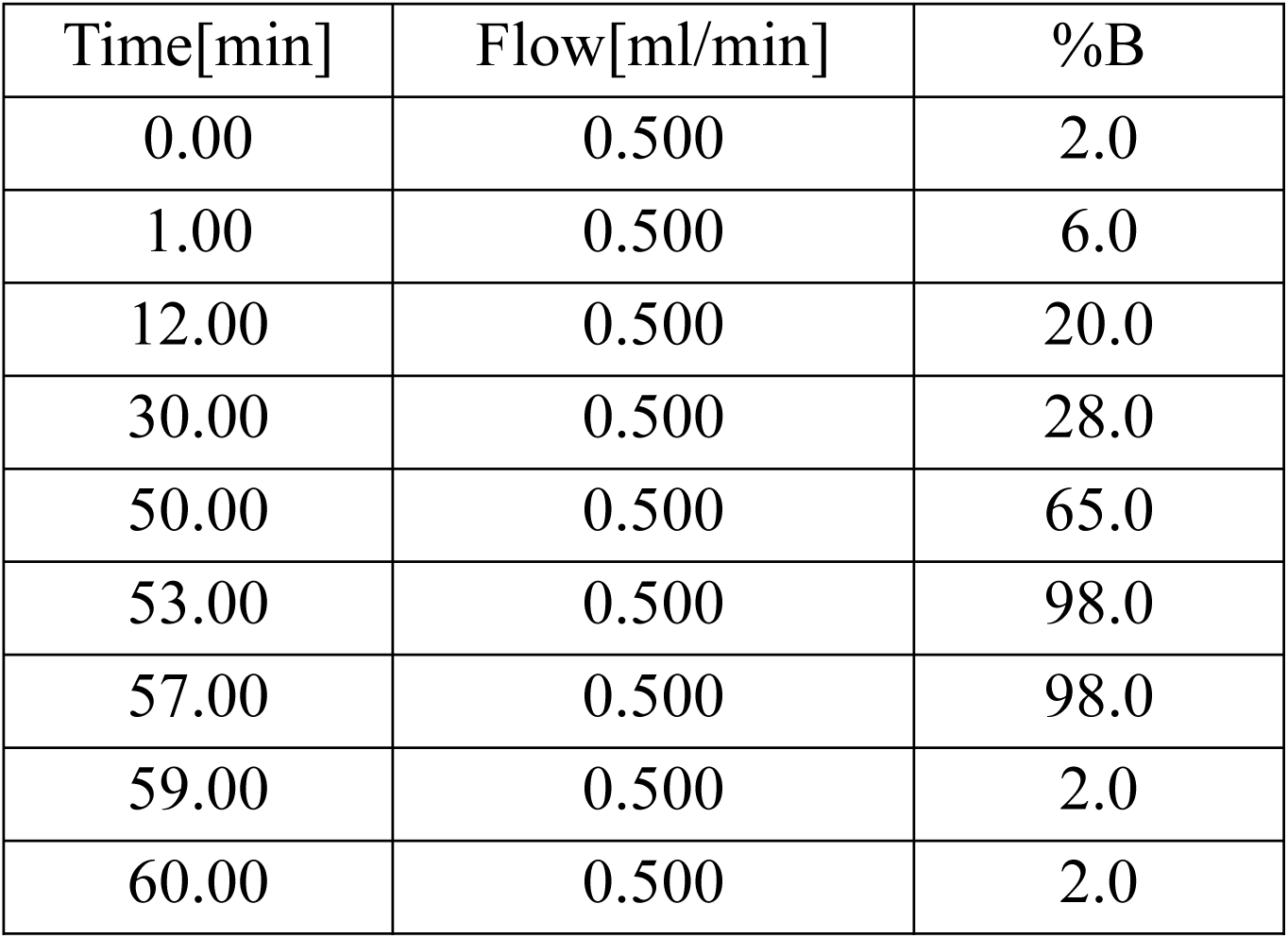
High pH Reverse Phase HPLC Fractionation Gradient Information.

| Time[min] | Flow[ml/min] | %B |
| --- | --- | --- |
| 0.00 | 0.500 | 2.0 |
| 1.00 | 0.500 | 6.0 |
| 12.00 | 0.500 | 20.0 |
| 30.00 | 0.500 | 28.0 |
| 50.00 | 0.500 | 65.0 |
| 53.00 | 0.500 | 98.0 |
| 57.00 | 0.500 | 98.0 |
| 59.00 | 0.500 | 2.0 |
| 60.00 | 0.500 | 2.0 |

**Supplementary Table 7.**
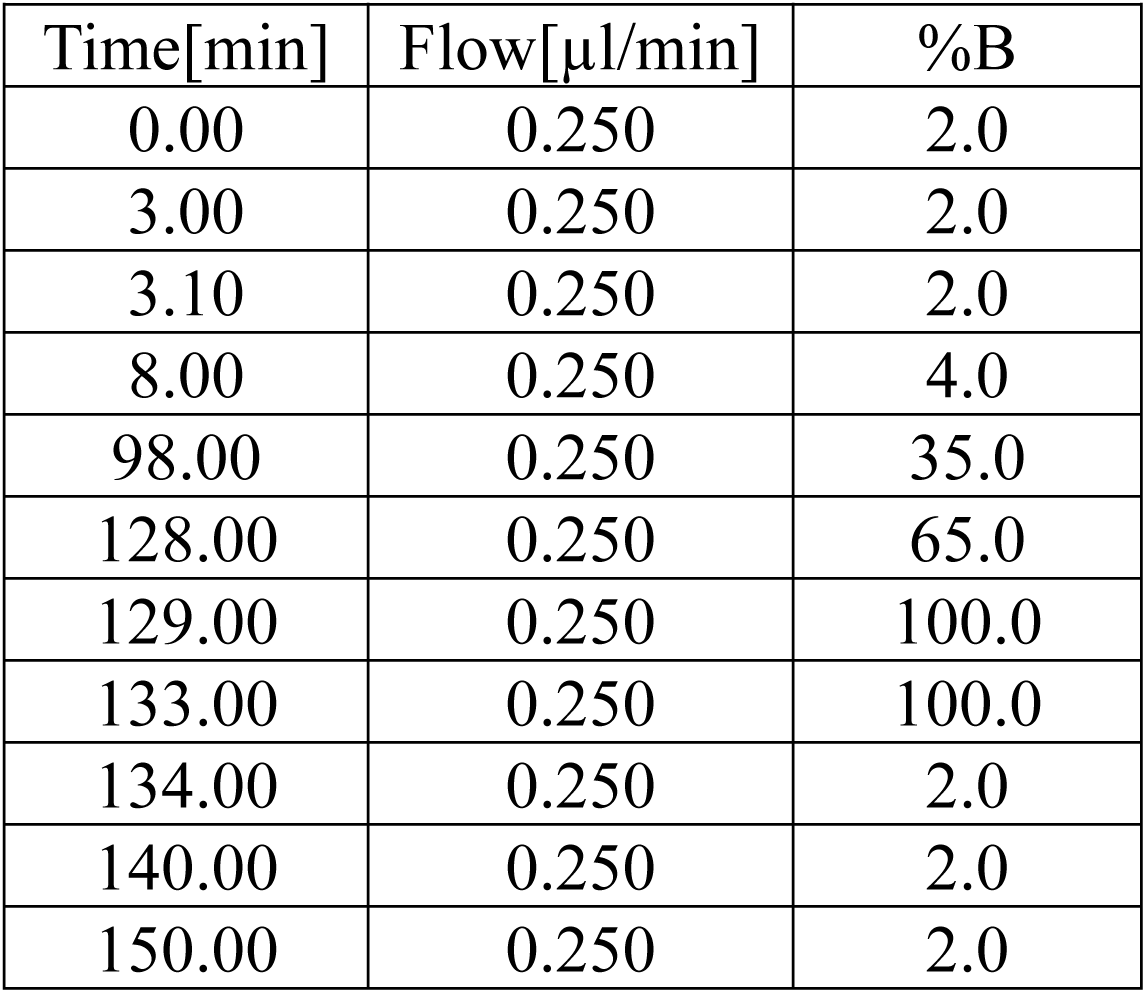
nanoLC-MS Gradient Information.

| Time[min] | Flow[μl/min] | %B |
| --- | --- | --- |
| 0.00 | 0.250 | 2.0 |
| 3.00 | 0.250 | 2.0 |
| 3.10 | 0.250 | 2.0 |
| 8.00 | 0.250 | 4.0 |
| 98.00 | 0.250 | 35.0 |
| 128.00 | 0.250 | 65.0 |
| 129.00 | 0.250 | 100.0 |
| 133.00 | 0.250 | 100.0 |
| 134.00 | 0.250 | 2.0 |
| 140.00 | 0.250 | 2.0 |
| 150.00 | 0.250 | 2.0 |

**Supplementary Table 8.** DIA segment 1 Precursor Scan Range Information.

| Segment 1<br>(400 – 800 m/z, IW 15 m/z, Overlap 1 m/z) |  |
| --- | --- |
| 399.5 – 415.5 | 609.5 – 625.5 |
| 414.5 – 430.5 | 624.5 – 640.5 |
| 429.5 – 445.5 | 639.5 – 655.5 |
| 444.5 – 460.5 | 654.5 – 670.5 |
| 459.5 – 475.5 | 669.5 – 685.5 |
| 474.5 – 490.5 | 684.5 – 700.5 |
| 489.5 – 505.5 | 699.5 – 715.5 |
| 504.5 – 520.5 | 714.5 – 730.5 |
| 519.5 – 535.5 | 729.5 – 745.5 |
| 534.5 – 550.5 | 744.5 – 760.5 |
| 549.5 – 565.5 | 759.5 – 775.5 |
| 564.5 – 580.5 | 774.5 – 790.5 |
| 579.5 – 595.5 | 789.5 – 800.5 |
| 594.5 – 610.5 |  |

**Supplementary Table 9.** DIA segment 2 Precursor Scan Range Information.

| Segment 2<br>(800 – 1200 m/z, IW 25 m/z, Overlap 1 m/z) |  |
| --- | --- |
| 799.5 – 825.5 | 999.5 – 1025.5 |
| 824.5 – 850.5 | 1024.5 – 1050.5 |
| 849.5 – 875.5 | 1049.5 – 1075.5 |
| 874.5 – 900.5 | 1074.5 – 1100.5 |
| 899.5 – 925.5 | 1099.5 – 1125.5 |
| 924.5 – 950.5 | 1024.5 – 1150.5 |
| 949.5 – 975.5 | 1049.5 – 1175.5 |
| 974.5 – 1000.5 | 1074.5 – 1200.5 |

