## Supplemental Tables for "Pharmaco-Pheno-Multiomic Integration Reveals Biomarker Profiles and Therapeutic Response Prediction Models in Leukemia and Ovarian Cancer"

**Supplementary Legends and Tables**

**Supplementary Table 1:** Clinical and Molecular overview of the AML patient cohort and tumor samples collected.

| **Model** | **FAB classification/WHO subtype** | **Cytogenetics** | **% blasts (peripheral blood)** | **White blood cell count (x10^9^/L)** | **Platelet count (x10^3^/ml)** | **CD34 status** | **IDH1/2 status** | **FLT3 status** | **NPM status** | **Diagnosis** | **Treatment history** | **Age** | **Gender** | **Ethnicity** |
| --- | --- | --- | --- | --- | --- | --- | --- | --- | --- | --- | --- | --- | --- | --- |
| CTG-2227 | M4 (myelomonocytic) | Not available | 89 | 288 | 51 | CD34+ | IDH1 mutant (R132H) | ITD mutant | mut | Relapsed | Pretreated | 59 | F | Caucasian |
| CTG-2229 | M1 (without maturation) | 46, XY, del(2)(p13p23), t(4;13)(q31;q34), add(4)(q25), del(6)(q13q25), t(9;22)(q34;q11.2), del(10)(q24), add(16)(q24) [20] | 95 | 134 | 11 | CD34+ bright | IDH1 mutant (R132C) | Wild type | Wild type | Refractory | Pretreated | 53 | M | Caucasian |
| CTG-2232 | NOS | Normal | Not available | 431 | 0 | Not available | Wild type | ITD mutant | mut | De novo | Naïve | 54 | F | Caucasian |
| CTG-2233 | AML-MLD; prior MDS/MPN | 47, XX, inv(10)(p11.2q21.2)c, +13 [6]/46, XX, inv(10)(p11.2q21.2)c [14] | 21 | 128 | 25 | CD34+ small subset/variable | IDH2 mutant (R140Q) | ITD mutant | Wild type | Secondary | Pretreated | 75 | F | Not available |
| CTG-2234 | NOS | Normal | 92 | 96 | 38 | CD34+ | Wild type | ITD mutant | mut | De novo | Naïve | 39 | M | Caucasian |
| CTG-2235 | AML-MLD; prior MPN | 46, XY, del(20)(q11.2q13.1) [20] | 89 | 155 | 43 | CD34+ | Wild type | Wild type | Wild type | Secondary | Pretreated | 66 | M | Caucasian |
| CTG-2236 | Biphenotypic | 46, XY, der(7)t(7;13)(q22;q13) [13]/48, idem, +9, +13 [7]/47, XY, +12 [1]/48, XY, +12, +13 [1] | Not available | 246 | Not available | CD34+ bright | Not available | ITD mutant | Wild type | Not available | Naïve | 75 | M | Not available |
| CTG-2238 | M4 (myelomonocytic) | Normal | 89 | 180 | 48 | CD34+ | Wild type | ITD mutant | Wild type | De novo | Naïve | 55 | M | Caucasian |
| CTG-2239 | NOS | Normal | 96 | 264 | 79 | CD34+ small subset/variable | Wild type | Not available | mut | De novo | Naïve | 79 | M | Black or African American |
| CTG-2240 | AML (11q23 abnormalities) | 46, XY, t(9;11)(p22;q23), add(10)(q24) [5]/46, XY [5] | 91 | 109 | 99 | CD34- | Wild type | Wild type | Wild type | De novo | Naïve | 75 | M | Not available |
| CTG-2453 | AML-MLD; prior MDS | Normal | 89 | 162 | 37 | CD34+ | Wild type | Non-ITD mutant (V579A) | mut | Secondary; refractory | Pretreated | Not available | M | Not available |
| CTG-2456 | NOS | 46, XX, del(7)(q22q36) [15] | 75 | 179 | 393 | CD34+ | Wild type | Wild type | Wild type | De novo | Pretreated | Not available | F | Caucasian |
| CTG-2457 | NOS | 44, XY, der(3)t(3;11)(p21;q13), del(5)(q?15q?34), del(6)(q12), der(7)t(6;7)(q12;p15), -11, der(17)t(11;17)(p11.2;p11.2), -18 [11]/44, idem, -del(6)(q12), +i(6)(p10) [5]/44, idem, -Y, +i(Y)(q10), -del(6)(q12), +i(6)(p10) [4] | 60 | 97 | 51 | CD34+ | Wild type | Wild type | Wild type | De novo | Naïve | Not available | M | Caucasian |
| CTG-2700 | M2 (with maturation) | 45, X, -Y [5]/46, XY [3] | 85 | 96 | 45 | Not available | Not available | Wild type | mut | De novo | Pretreated | Not available | M | Caucasian |
| CTG-2701 | M5 (monocytic; M5a and M5b) | Normal | 40 | 166 | 29 | CD34+ dim | Not available | ITD mutant | mut | De novo | Naïve | Not available | M | Caucasian |
| CTG-2702 | M4 (myelomonocytic) | 46, X, -Y, +8 [13]/46, XY [2] | 38 | 149 | 505 | Not available | Not available | Wild type | N/A | Relapsed | Pretreated | Not available | M | Caucasian |
| CTG-2704 | M1 (without maturation) | Normal | 82 | 143 | 33 | CD34- | Not available | ITD mutant | N/A | De novo | Naïve | Not available | F | Caucasian |
| CTG-2774 | Not available | Normal | 13.6 | 64 | 106 | CD34- | Wild type | Wild type | N/A | Recurrent | Pretreated | 65 | M | Caucasian |
| CTG-2775 | AML-MLD; prior MPN | 47, XY, +6 [6]/46XY [15] | 31.5 | 39 | 16 | CD34+ partial/variable | Not available | Not available | N/A | Secondary | Pretreated | 64 | M | Not available |
| CTG-3438 | M4 (myelomonocytic) | Normal | 66 | 50 | 20 | CD34+ | Wild type | Wild type | Wild type | Relapsed | Pretreated | 62 | F | Caucasian |
| CTG-3439 | Not specified (prior MDS) | Normal | 67 | 218 | 55 | CD34+ variable | Wild type | ITD mutant | Wild type | Secondary | Pretreated | 64 | M | Caucasian |
| CTG-3441 | M1 (without maturation) | Not available | 96 | 186 | N/A | CD34- | Wild type | ITD mutant | mut | De novo | Pretreated | 80 | M | Caucasian |
| CTG-3679 | M0 (minimally differentiated) | 46, XX, i(17)(q10) [14]/46, XX [6] | 71 | 256 | N/A | CD34+ | Wild type | Non-ITD mutant (T582_E598dup) | Wild type | De novo | Naïve | 54 | F | Caucasian |

**Supplementary Table 5.** Clinical and Molecular overview of the Ovarian patient cohort and PDX Models Established.

| **Model** | **Tumor status** | **Harvest site** | **Histology** | **Tumor grade** | **Diagnosis** | **Treatment history** | **Disease stage** | **Age** | **Ethnicity** |
| --- | --- | --- | --- | --- | --- | --- | --- | --- | --- |
| CTG-0258 | Metastatic | Small bowel | Papillary carcinoma, serous origin | Poorly differentiated | N/A | N/A | N/A | N/A | N/A |
| CTG-0486 | Metastatic | Small bowel | Papillary carcinoma, serous origin | Poorly differentiated | N/A | N/A | III | 77 | N/A |
| CTG-0703 | Metastatic | Abdomen | Serous carcinoma | Poorly differentiated | First diagnosis | Naive | III | 59 | Asian |
| CTG-0712 | Local metastatic | Omentum | Papillary carcinoma, serous origin | Poorly differentiated | N/A | Pretreated | IV | 72 | Caucasian |
| CTG-0791 | Metastatic | Abdomen | Papillary carcinoma, serous origin | Poorly differentiated | First diagnosis | Pretreated | III | 46 | Caucasian |
| CTG-0868 | Metastatic | Liver | Endometrioid carcinoma | Poorly differentiated | Recurrent | Pretreated | IV | 72 | Caucasian |
| CTG-0897 | Metastatic | Diaphragm | Serous carcinoma | Poorly differentiated | Recurrent | Pretreated | III | 62 | Caucasian |
| CTG-0947 | Metastatic | Lymph node | Epithelial | Poorly differentiated | Recurrent | Pretreated | III | 59 | Caucasian |
| CTG-0956 | Metastatic | Pelvis | Serous carcinoma | Well differentiated | Recurrent | Pretreated | IV | 73 | Caucasian |
| CTG-0958 | Metastatic | Uterine adnexa | Papillary carcinoma, serous origin | Poorly differentiated | Recurrent | Pretreated | IV | 62 | Caucasian |
| CTG-0964 | Metastatic | Peritoneum | Papillary carcinoma, serous origin | Poorly differentiated | Recurrent | Pretreated | IV | 56 | Asian |
| CTG-0992 | Metastatic | Liver | Carcinoma | Poorly differentiated | Recurrent | Pretreated | III | 71 | Caucasian |
| CTG-1086 | Metastatic | Lymph node | Papillary carcinoma, serous origin | Poorly differentiated | Recurrent | Pretreated | III | 65 | Caucasian |
| CTG-1166 | Metastatic | Neck | Papillary carcinoma, serous origin | Poorly differentiated | Recurrent | Pretreated | IV | 60 | Caucasian |
| CTG-1180 | Metastatic | Liver | Endometrioid carcinoma | Poorly differentiated | Recurrent | Pretreated | IV | 73 | Caucasian |
| CTG-1305 | Primary | Ovary | Serous cyst-adenocarcinoma | Undifferentiated | First diagnosis | Naive | III | 60 | Caucasian |
| CTG-1423 | Primary | Ovary | Mixed epithelial carcinoma | Poorly differentiated | First diagnosis | Naive | II | 46 | Asian |
| CTG-1498 | Metastatic | Lymph node | Carcinoma | Poorly differentiated | Recurrent | Pretreated | III | 55 | Asian |
| CTG-1602 | Local metastatic | Uterus | Dedifferentiated carcinoma | Poorly differentiated | N/A | N/A | N/A | 53 | Not available |
| CTG-1703 | Local metastatic | Pelvis | Serous carcinoma | Poorly differentiated | Recurrent | Pretreated | III | 73 | Not available |
| CTG-2213 | Metastatic | Abdomen | Clear cell carcinoma | Poorly differentiated | First diagnosis | Naive | III | 55 | Caucasian |
| CTG-3226 | Metastatic | Lymph node | Carcinoma | Poorly differentiated | Recurrent | Pretreated | IV | 56 | Caucasian |
| CTG-3383 | Metastatic | Lymph node | Serous carcinoma | Undifferentiated | Recurrent | Pretreated | IV | 57 | Caucasian |
